# Efficacy of DNA Methyltransferase Inhibitor Immune Priming Therapy in Combination with PD-1 Inhibitors to Treat High-Risk Pediatric Brain Tumors

**DOI:** 10.1101/2025.05.14.654106

**Authors:** Deepak Kumar Mishra, Shelli M. Morris, Dean Popovski, Emily J. Girard, Andrew Bondoc, Shiva Senthil Kumar, Augusto Faria Andrade, Xiaoting Zhu, Fupan Yao, Mi-Youn Brusniak, Umaru Banlanjo, Erin E. Crotty, Ken Brasel, Fiona Pakiam, Caterina Russo, Michele Zeinieh, Matt C. Biery, Margo Coxon, Heather Conti, Midori Clarke, Mei Lu, James Rutka, Dhana Llivichuzhca-Loja, Liza Konnikova, Maryam Fouladi, Nada Jabado, Annie Huang, James M. Olson, Rachid Drissi

## Abstract

**Background:** Despite intensive therapies, outcomes for high-risk pediatric brain tumors (PBTs) remain dismal, prompting the search for novel treatments. DNA methyltransferase inhibitors (DNMTi) have been shown to prime tumors to improve response to checkpoint inhibition. The aim of this study was to investigate the potential of decitabine (DAC), in combination with a PD-1 inhibitor, to improve survival in pediatric high-risk brain tumor models.

**Methods:** Analysis of human PBT datasets was performed to determine gene expression levels of immune cell associated markers. Tumor response to DAC, with or without a PD-1 inhibitor, was tested in murine models representing H3-wildtype diffuse intrinsic pontine glioma (DIPG), H3K27-mutant diffuse midline glioma (DMG), atypical teratoid rhabdoid tumor (ATRT), and medulloblastoma (MB). CyTOF analysis of allograft tumors was performed to characterize changes within the tumor microenvironment.

**Results:** Analysis of PBT subtypes revealed heterogeneous expression of immune cell markers, checkpoint receptors, and MHC molecules. DAC treatment decreased DNA methylation and increased neoantigen expression in human and mouse tumor cells. DAC alone or in combination with a PD-1 inhibitor resulted in prolonged survival in syngeneic mouse models of DIPG and ATRT but not DMG and MB models. CyTOF analysis of mouse tumors revealed changes in local immune cell infiltration upon combination treatment.

**Conclusions:** DAC in combination with a PD-1 inhibitor can alter the immune microenvironment in mouse tumor models. Changes were observed in H3-wildtype DIPG and ATRT models, suggesting that certain tumor subtypes may respond to checkpoint blockade after immune augmentation with DNMTi.

**Key Points:**

1. PBTs show heterogenous expression of immune cell infiltrates
2. DAC or DAC plus a PD-1 inhibitor shows extension of survival in H3-wildtype DIPG and ATRT mouse models
3. Myeloid-derived suppressor cell abundance could be a major contributing factor to treatment response

**Importance of the Study:** Children with high-risk PBTs face dismal outcomes. Immune checkpoint inhibitor (ICI) successes have been demonstrated in a variety of adult malignancies; however, such beneficial outcomes have not been realized in PBTs. Here we investigate single and combination treatment of DNMTi and PD-1 checkpoint inhibition in syngeneic mouse models of high-risk PBTs. Our results suggest that some H3-wildtype DIPG and ATRT tumor types may be responsive to checkpoint therapy post immunomodulation and warrant further investigation.

## INTRODUCTION

Central nervous system (CNS) tumors remain the leading cause of cancer-related deaths in children.^1^ Clinical outcomes for poor prognosis pediatric brain tumors (PBTs), such as diffuse midline glioma (DMG)^2^, including diffuse intrinsic pontine glioma (DIPG), high-grade glioma (HGG)^3^, atypical teratoid rhabdoid tumor (ATRT)^4^, and *MYC* and *MYCN*-amplified medulloblastoma (MB)^5^, remain poor. Multimodal therapy that consists of surgery, radiotherapy (RT) ± chemotherapy has yielded 5-year event-free survival (EFS) of 0-50% in this population. Unfortunately, survivors experience significant treatment-related sequelae, including neurocognitive and neuroendocrine dysfunction, ototoxicity, and secondary malignancies that have long-term impacts on patients, their families, and the healthcare system.^6^ To address this clinical deficit there is an urgent need to develop and implement novel treatment paradigms that improve overall survival outcomes and reduce the morbidity associated with current therapies.

Immune checkpoint inhibitors (ICIs), such as PD-1, PD-L1, and CTLA-4 inhibitors, have revolutionized cancer treatment in recent years and have demonstrated profound clinical survival benefit in subsets of adult tumors.^7,8^ Moreover, emerging evidence indicates that conventional (chemotherapy, RT) and molecularly targeted therapies can have immunomodulatory effects beyond cell-intrinsic and genetic mechanisms.^9,10^ These effects on the immune system are thought to be due, in part, to increasing the expression of tumor associated antigens and promoting lymphocytic infiltration in a process that can turn an immunologically inert tumor into an active one. As a result of their success in adult tumors, ICIs are thought to provide an attractive opportunity to target PBTs with a child’s own immune system, reducing the morbidity of surgical resection, RT, and chemotherapy in young patients and potential disruption of neuronal maturation. Unfortunately, early results of pediatric trials with single agent immune checkpoint blockers have been, with rare exceptions, disappointing.^11–15^ Therefore, it is likely that a combinatorial treatment with other agents may be necessary if ICIs are to play a role in PBT therapy.

One challenge with ICI therapy in PBTs is the lower mutational burden as compared to adult tumors, resulting in fewer neoantigens that can trigger an immunological response.^16^ Furthermore, downregulation of MHC expression on tumor cells, leading to poor antigen presentation, has been reported as a form of immune evasion from cytotoxic CD8 T cells.^17^ Additional factors like cytokines, chemokines, and immunosuppressive cells, including regulatory T cells (Tregs) and myeloid-derived suppressor cells (MDSCs), can influence the tumor immune microenvironment towards an inert ‘cold’ state. Thus, intricate immune evasion mechanisms can lead to reduced lymphocytic infiltrate within the tumor microenvironment (TME) and compound the reduced immunological response to ICIs.^18^

Immune priming represents a promising strategy to augment the response to ICIs in PBTs. DNA methyltransferase inhibitors (DNMTi), such as azacitidine (AZA) and 5-aza-deoxycytidine (decitabine, DAC), are such agents that have been shown in other tumor types to promote an immune priming response by inducing neoantigen expression through global DNA hypomethylation.^16,19,20^ Preclinical studies of myelodysplastic syndromes and gliomas have demonstrated DAC’s role as an immunomodulator through increased T lymphocyte infliltration.^21–24^ In a murine tumor model expressing human papillomavirus 16, treatment with DAC along with a histone deacetylase inhibitor, triggered MHC I expression, epigenetic activation of antigen-processing, and increased susceptibility to cytotoxic T cell lysis.^25^ With this information in mind, and to address the lack of response to ICIs in pediatric patients, we evaluated the effect of DNMTi on human and mouse PBT tumor models and investigated whether the combination of DAC treatment and PD-1 checkpoint inhibition could increase survival in multiple syngeneic mouse models representing various molecular types of PBTs.

## MATERIALS AND METHODS

### Expression Data

The DIPG data set included 28 patients’ paired tumor and normal tissues from the frontal lobe. RNA-seq data were used for the gene expression analysis. Tumor samples included 20 with H3.3K27M mutation, 7 with H3.1K27M mutation, and one histone WT. The data were processed following the same process as described.^26^ Gene expression was represented by normalized gene counts, and Wilcoxon Rank Sum test was used to compare the difference between tumor and normal tissues. P<0.05 was considered significant.

Primary ATRTs were downloaded and processed on the Illumina HT-12 platform as described.^27^ A total of 58 samples were processed, including 21 fetal brain samples, 15 SHH, 14 TYR, and 8 MYC primary human tumor samples. Primary ATRTs and fetal brain samples were batch corrected by ComBat.^28^ Differential testing was performed by applying Wilcoxon rank-sum test against fetal brain as control.

The gene expression data of medulloblastoma were obtained from the R2:Genomics Analysis and Visualization platform (https://r2.amc.nl). The Swartling data set (GSE124814) was analyzed.^29^ This set contained 23 integrated transcription datasets that were batch corrected and included 291 normal brain, 118 WNT, 405 SHH, 233 group 3, 530 group 4 samples. The Cavalli data set of 763 primary samples (GSE85217) including 70 WNT, 223 SHH, 144 group 3, and 326 group 4 was also analyzed.^30^ The gene expression level of each marker was compared by ANOVA test. Stars represent the significance of comparing each group against base-mean.

CIBERSORT was performed using the CIBERSORTx online tool (https://cibersortx.stanford.edu/).^31^ Gene expression data from the DIPG, ATRT and medulloblastoma data sets were input and the abundance of immune cell types was estimated using the “Impute Cell Fractions” function and the LM22 (22 immune cell types) dataset as the signature matrix file. Default parameters were used as disable quantile normalization, run in absolute mode, using 1000 permutations for significance analysis.

### Cell Lines

See Supplemental Materials for a complete list of both human and mouse cell lines.^32–42^

### *In Vitro* Drug Treatment

Human and mouse cell lines were plated and allowed to adhere overnight. Cells were incubated with or without various concentrations of DAC or AZA (Decitabine; Azacitidine, Selleckchem) for either 24, 36, or 48 hours, with daily media drug changes. A positive control utilizing 50 ng/mL of human or murine interferon-gamma (IFNγ, Peprotech) was included. At the end of the treatment period, cells were collected, counted, and analyzed for viability via Trypan blue staining.

### Flow Cytometry

Cultured cells were trypsinized and single cell suspensions were incubated with human Fc block (TruStain FcX, BioLegend), stained according to standard protocols, and run on an Agilent NovoCyte Flow Cytometer. Data were analyzed with NovoExpress Software and FlowJo Software version 10. See Supplemental Materials for list of antibodies.

### Quantitative RT-PCR Analysis of Neoantigens

Cultured cells were treated with 0.1 µM, 0.5 µM, or 1 µM DAC for 3 days. Total RNA extraction was performed from cell lysates using TRIzol (Invitrogen) and the Qiagen RNeasy Mini Kit (Qiagen) and quantified by Nanodrop. cDNA was synthesized using the High-Capacity RNA-to-cDNA Kit (Applied Biosystems). RT-qPCR was performed with an ABI QuantStudio 5 Real-Time PCR System (ThermoFisher) using TaqMan Gene Expression Universal Master Mix (Applied Biosystems) and TaqMan Gene Expression primers (ThermoFisher). A total of 40 cycles was performed. For those neoantigens that were undetectable, a Ct value of 40 was used to calculate ΔΔCt and RNA fold change. See Supplemental Materials for list of TaqMan assays.

### Dot Blot Assay for the Evaluation of DNA Methylation

For cell lines and tissues, genomic DNA (gDNA) was isolated using Qiagen gDNA isolation kit, denatured in 0.1 M NaOH for 10 min at 95°C, neutralized with 1 M NH_4_OAc on ice, diluted two-fold using SSC 2x buffer and 200 μL of the serial diluted gDNA was spotted on nitrocellulose membrane followed by UV crosslinking. The membrane was blocked by soaking in 5% BSA in TBS-T for 1 hr followed by incubation with a mouse anti-5-methylcytosine (5-mC) monoclonal antibody (Sigma-Aldrich, 1:1,000) in TBS-T at 4 °C overnight. The next day, the membrane was washed 3x for 5 min each in TBS-T, and then incubated with a secondary antibody, HRP-conjugated anti-mouse immunoglobulin-G (IgG) (1:5,000) in TBS-T for 1 hr at room temperature. Following 3x 5 min washes, the membrane was imaged using chemiluminescence. As a loading control, the membrane was re-stained using methylene blue.

### Animal Research and Syngeneic Tumor Models

All mouse experiments were conducted in accordance with the National Institutes of Health (NIH) *Guide for the Care and Use of Laboratory Animals*, with approval from the Institutional Care and Use Committees at each site. Female C57BL/6J mice (The Jackson Laboratory), were used when 4-8 weeks old. All mice were group housed with unrestricted mobility and free access to food and water. IUE-24-C5 flank tumors were established by implanting 500,000 cells. Syngeneic tumors were surgically implanted in the pons [DIPG model IUE-24-C5 (Pdgfra^D842V^, dominant neg *Trp53*, H3 WT), 100,000 cells]^32^, [DMG model IUE-K27M-APP (H3.3K27M, *Trp53*-*Atrx*-KD, PDGFRA-overexpression), 150,000 cells]^34,43^, cerebral cortex [ATRT model SU2C_54_i_5 (ROSA-CreER/SNF5^fl/fl^), 1×10^6^ cells], or cerebellum [medulloblastoma models SHH57835 (*Ptch1*^+/-^, *Trp53* loss), 50,000 cells; 7444 (*c-Myc* CRISPR, *Trp53* loss, *Cdkn2c* loss), 50,000 cells]^33,35^ of mice as previously described with tumor cells originating from symptomatic intracranial tumors in donor mice or from tissue culture.^44^ The time between surgical implantation of tumors and initiation of treatment was based on previously established growth latency in each model. Mice were randomly assigned to the following treatment groups and monitored daily for symptoms of tumor burden. Treatment Groups: 1-Vehicle + Isotype; 2-DAC + Isotype; 3-Vehicle + 4H2; 4-DAC + 4H2. DAC was dosed intraperitoneal (IP) Monday, Tuesday, and Wednesday of each week at 0.5 mg/kg. Isotype control and 4H2 (an anti-mouse PD-1 antibody kindly provided by Bristol Myers Squibb) were dosed IP at 200 μg per mouse on Monday and Friday of each week, beginning on the Friday of the first week of dosing. Mice were euthanized according to pre-determined humane endpoint criteria including when tumor burden began to impede normal behavior and functioning and/or 20% body weight loss occurred.

### Tumor Imaging

Mice implanted with IUE-24-C5 tumors were administered 200 μL of 3 mg D-Luciferin/20 g mouse weight (RediJect D-Luciferin Ultra, PerkinElmer) by IP injection and anesthetized by isoflurane inhalation after 2 minutes. Luminescence was measured in anesthetized mice on a Xenogen IVIS instrument with 1 min exposure and quantified using LivingImage software (version 4.0, Revity). MRI imaging was used to confirm tumor formation and growth in ATRT mice. Briefly, mice were anaesthetized using isoflurane (maintained on 1-1.5 %), placed in the MRI machine with ongoing vital monitoring, The contrast agent utilized was 50 mg/kg manganese chloride (MnCl_2_) administered IP, 24 hours prior to imaging. Mice were imaged once per week, every three weeks, with a T1-weighted M contrast-enhanced image at 75 μm isotropic resolution using a Bruker 7T BioSpec 70/30 USR MRI machine.

### Immunohistochemistry and Image Quantification

Refer to supplemental materials for complete list of antibodies. Digital slides were created using Aperio FL and uploaded to the HALO Image Analysis Platform (Indica Labs) for immune cell quantification. Immune cells were detected by positive staining within whole brain sections as a percentage of total tissue area (µm^2^) and analyzed by CytoNuclear or Immune Cell modules (Indica Labs).

### Cytometry by Time of Flight (CyTOF)

Tumor samples were collected prior to experimental endpoint (DIPG, ATRT, DMG models) or following one week of dosing (MB) and digested utilizing the Miltenyi Biotec Brain Tumor Dissociation Kit into single cell suspensions per the manufacturer’s instructions. Single cell suspensions underwent CyTOF staining per previously published protocol.^45^ Briefly, cells were washed with cell-staining buffer (CSB): DPBS with 0.5 % bovine serum albumin (Sigma) and 0.02 % sodium azide, and then incubated with Human TruStain FcX (BioLegend). Cell viability was assessed with 103Rh (Fluidigm). After washing with CSB, cells were stained with a surface staining antibody cocktail. For intracellular staining, cells were washed with CSB and incubated in FoxP3 fixation and permeabilization (Invitrogen). After permeabilization, cells were washed with FoxP3 wash buffer (Invitrogen) and stained with an intracellular antibody cocktail. After staining and washing, cells were fixed with 1.6 % paraformaldehyde (Sigma) and stored overnight in CSB at 4°C. On the following day, cells were labeled with 191Ir/193Ir DNA Intercalator (Fluidigm) and then washed with CSB. On the day of analysis, cells were washed with MilliQ water and resuspended in EQ Beads (Fluidigm) diluted 1:10. Analysis was done on a Fluidigm mass cytometer. Data were exported as FCS files.

FCS files were uploaded to Premium Cytobank (Beckman Coulter) where they were manually gated for events that were positive for both DNA dyes (Ir191/Ir193), negative for 103Rh for viable cells, singlets based on event length, negative for 140Ce to remove bead signals, and positive for CD45^+^. These events were exported as FCS files and imported into cytofkit2.^46^ The data loading settings were chosen as follows: merge method ceil, fixed number of cells from each FCS file, and transformation method cytofAsinh. Dimensionality reduction was performed with t-distributed stochastic neighbor embedding (t-SNE) and cell clustering was carried out with Rphenograph with a k value of 30. Clusters were manually annotated using marker expression heatmaps. Cluster abundance data and mean metal intensity (MMI) values were used to generate plots utilizing GraphPad Prism 9.

## RESULTS

### Pediatric brain tumors have heterogenous expression of immune associated markers

To determine the initial immunological state of PBTs, we evaluated immunological markers in patient-derived tumor tissues using RNA-seq or microarray-based gene expression analyses **(Figure 1, Supplementary Figures 1-4)**. Markers focused on human data sets that were compatible with our downstream CyTOF analyses of mouse preclinical brain tumor models **(Figures 5 and 6)**. For DIPG samples, we compared the expression data generated using RNA-seq from a cohort of patient tumors to matched normal frontal lobe tissue collected at autopsy.^26^ DIPG tumors are located in the pons and diagnosed using MRI. On the other hand, DMGs are diagnosed by biopsy and have a specific mutation in the *H3-3A* gene (H3K27M mutant). This data set contained both H3K27M mutants and a histone wildtype sample. While the marker of proliferation, *MKI67,* was increased in tumor samples, DIPG tumor samples showed no significant upregulation of immune checkpoint markers *PD-L1 (CD274)* and *CTLA-4* compared to matched normal tissue **(Supplementary Figure 1)**. However, there were some individuals who exhibited either high or low expression of *PD-L1* and/or *CTLA-4*. ATRT tumors can be divided into 3 molecular subtypes – TYR, SHH, and MYC. Like the DIPG patient samples, ATRT tumors had significantly increased *MKI67* expression, but only the TYR subgroup exhibited an increase in *PD-L1* while the TYR and MYC subgroups had significant increases in *CTLA-4* levels **(Supplementary Figure 2)**. Medulloblastomas are divided into 4 different subtypes – Sonic Hedgehog (SHH), WNT, Group 3 and Group 4. For medulloblastomas, expression analysis from two different data sets, representing 4 different subtypes, exhibited variation in expression of *PD-L1* and *CTLA-4* among subtypes of the disease. In general, *MKI67* was increased in most tumor types, with WNT tumors exhibiting the biggest increase. *PD-L1* expression was significantly increased in WNT samples while *CTLA-4* expression was found to be increased in the SHH subtype **(Supplementary Figures 3 and 4).**^47^ This heterogeneity of expression was also observed for MHC I and MHC II genes **(Supplementary Figures 1-4)** among the three disease types. We evaluated the expression of several immune-associated markers in these data sets. DIPG, ATRT, and SHH-activated medulloblastoma tumors had an increase in leukocytes compared to normal brain based on *PTPRC* (CD45) expression **(Supplementary Figures 1-4)**. Interestingly, we observed changes in various T cell markers **(Figure 1)**. Specifically, DIPG, ATRT (TYR, MYC), and medulloblastoma (SHH) had increased levels of CD4 expression, also known to be expressed by human monocytes and macrophages^48^, while CD8 expression tended to be decreased, except in the case of WNT-activated medulloblastoma. Myeloid-derived cells, as detected by *CD68* expression, were in general increased in DIPG, ATRT, and medulloblastoma tumor samples compared to normal brain. Interestingly, CD163, a marker associated with immunosuppressive myeloid cells, was also elevated in DIPG, ATRT, and medulloblastoma (WNT, SHH) tumors **(Supplementary Figures 1-4)**. Furthermore, the T cell stimulatory molecule *CD86* showed higher expression in tumors, while the immunosuppressive phenotypic marker *CCR2* expression varied among diseases. To estimate the relative ratio of the different immune populations in these data sets, CIBERSORT was performed **(Supplementary Figure 5)**. In general, DIPG, ATRT, and medulloblastoma (SHH) tumors contained increased number of cells with macrophage M2 gene signature.

**Figure 1.**
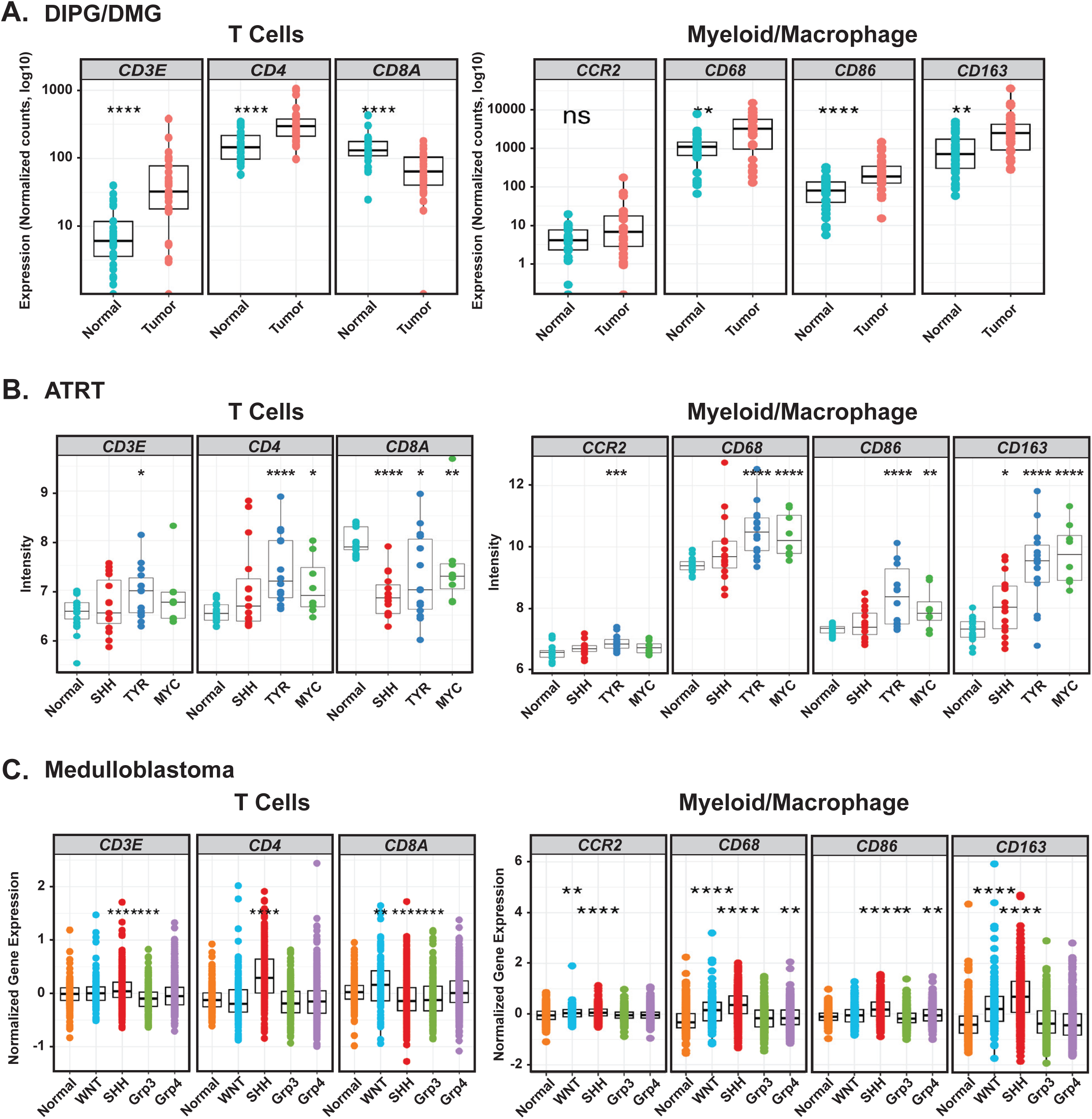
Pediatric brain tumors are heterogenous for select immune associated markers. (A) Expression analysis of 28 DIPG/DMG patients’ paired tumor and normal tissue. (B) Expression analysis of ATRT tumors including 15 SHH, 8 MYC, 14 TYR samples, plus 21 normal fetal brain samples. (C) Gene expression analysis of the Swartling data set which contains 23 integrated and batch corrected transcription datasets that include 291 normal brain, 118 WNT, 405 SHH, 233 group 3, 530 group 4 samples. ns = *P* > 0.05. Stars represent: * *P* ≤ 0.05, ** *P* ≤ 0.01, *** *P* ≤ 0.001, **** *P* ≤ 0.0001.

### Pediatric brain tumor cell lines exhibit heterogenous surface expression of MHC I and PD-L1

We evaluated the cell surface expression of MHC I and PD-L1 among different PBT cell lines using flow cytometry **(Figure 2A)**. MHC I expression was detected across multiple PBT cell lines. DMG/DIPG cells showed consistent expression of MHC I and variable expression of PD-L1. Medulloblastoma and ATRT lines were mixed for MHC I expression but tended to have lower baseline levels of PD-L1 on their surface. High-grade gliomas were quite variable in the expression of both surface proteins, suggesting heterogeneity with this tumor type, consistent with our molecular and genomic understanding of pHGG.

**Figure 2.**
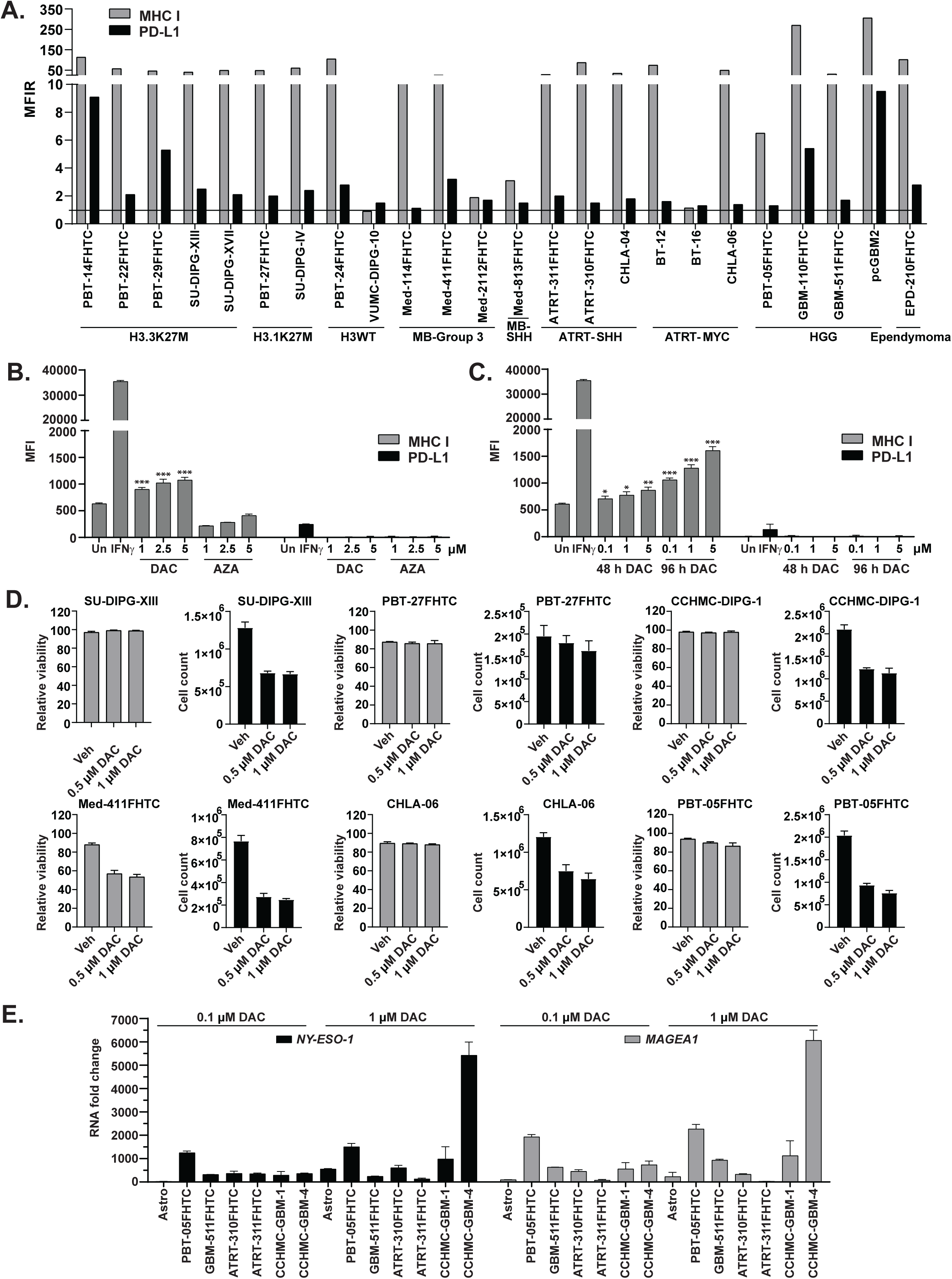
DNA methyltransferase inhibitor DAC induces MHC I and neoantigen expression in human pediatric brain tumor cells. (A) Baseline expression of MHC I and PD-L1 in a variety of pediatric brain tumor cell lines. MFIR equals the median fluorescence intensity of stained cells divided by the isotype control. Horizontal line indicates an MFIR of 1. (B) Induction of MHC I expression in PBT-05FHTC cells following a 2-day treatment of DAC or AZA. Un = Untreated. IFNγ (50 ng/ml for 48 hours) was used as a positive control. Stars represent: *P* = 0 - 0.001 ***, 0.001 - 0.01 **, 0.01 - 0.05 * (C) Changes in MHC I expression by DAC is dose and time dependent in PBT-05FHTC. (D) Cytotoxicity was evaluated after treating various cell lines with 0.5 µM, and 1 µM of DAC for 72 hours. (E) Quantitative PCR evaluation of *NY-ESO-1* and *MAGEA1* neoantigen gene expression in multiple PBT lines following 3-day DAC treatment. Astro, normal human astrocytes.

### DAC induces MHC I and neoantigen expression in pediatric brain tumor cells

Based on previous studies, DNMT inhibitors are known to reduce global DNA methylation^49^ and increase MHC I and neoantigen expression^20^, making the tumor cells more visible to the immune system.^50^ Therefore, we tested two FDA-approved DNMTi, azacitidine (AZA) and decitabine (DAC), in the pediatric high-grade glioma cell line, PBT-05FHTC, which expresses relatively low baseline levels of MHC I. Interferon-gamma (IFNɣ) was used as a positive control. We found DAC to be more effective in inducing MHC I expression as compared to AZA, which exhibited a slight decrease in MHC I, when dosed at the same concentrations **(Figure 2B)**. Furthermore, DAC treatment exhibited both dose and time-dependent induction of MHC I expression **(Figure 2C)** when tested on a patient-derived cell line at three concentrations (0.1, 1, and 5 µM) for 48 and 96 h. In contrast, no significant change in PD-L1 levels was observed. Additionally, DAC showed minimal cytotoxic effect (based on viability and cell counts) on most of the patient-derived cell lines except Med-411FHTC at both concentrations (0.5 and 1 µM) after a 3-day treatment **(Figure 2D).** We confirmed the induction of two known DAC inducible cancer testis antigens (CTA) in multiple PBT cell lines after 0.1 µM or 1 µM DAC treatment for 72 h **(Figure 2E)**. Cell lines showed variable induction of *NY-ESO-1* and *MAGEA1*, confirming that DAC treatment can induce expression of these neoantigens across a variety of brain tumor cell types.

### DAC influences cell viability, global methylation, and neoantigen expression in mouse syngeneic brain tumor cell lines

Prior to *in vivo* efficacy experiments in immune-competent mouse models, we confirmed the functional activity of DAC on mouse syngeneic cell lines representing a variety of PBT molecular subtypes. We focused on the DIPG model IUE-24-C5 (Pdgfra^D842V^, dominant negative *Trp53*, H3 WT),^32^ the DMG model IUE-K27M-APP (H3K27M, *Trp53-Atrx*-knockdown, PDGFRA-overexpression),^30,34^ and the medulloblastoma models SHH57835 (*Ptch1^+/-^*, *Trp53* loss) and 7444 (*c-Myc* CRISPR, *Trp53* loss, *Cdkn2c* loss).^33,35^ Initially, we conducted a 7-day *in vitro* test consisting of continuous DAC exposure on IUE-24-C5 cells and observed DACs cytotoxic effects **(Supplemental Figure 6)**. As a result, a shorter treatment plan akin to the human PBTs was selected for *in vitro* testing. The effect of DAC on cell viability and growth was tested at two concentrations (0.5 µM and 1 µM) for 72 h. Similar to the human cell lines, DAC showed primarily cytostatic effects at both concentrations **(Figure 3A)**. In addition, DAC treatment (0.5 µM) induced MHC I expression after 2- and 4-days treatment in a subset of mouse lines **(Supplementary Figure 7).** Global DNA methylation status of these cell lines using dot blot analysis showed significant reduction in DNA methylation after 72 h of treatment **(Figure 3B)**. Next, we confirmed the induction of CTA expression based on previously defined targets of decitabine in different mouse cell lines after 0.5 µM DAC treatment for 72 h **(Figure 3C)**^20,51^. Each cell line showed induction of CTA expression; however, there was variation among the lines suggesting that DAC treatment can affect cells differently. Medulloblastoma lines showed upregulation of *PrameL1* and *Trap1a*. The IUE-24-C5 DIPG cell line showed upregulation of *Trap1a* and *Rhox5* while the DMG line IUE-K27M-APP exhibited induction of *Trap1a*. Overall, these data show that DAC is also a potent epigenetic modulator of mouse syngeneic lines, similar to our observations in the human-derived PBTs.

**Figure 3.**
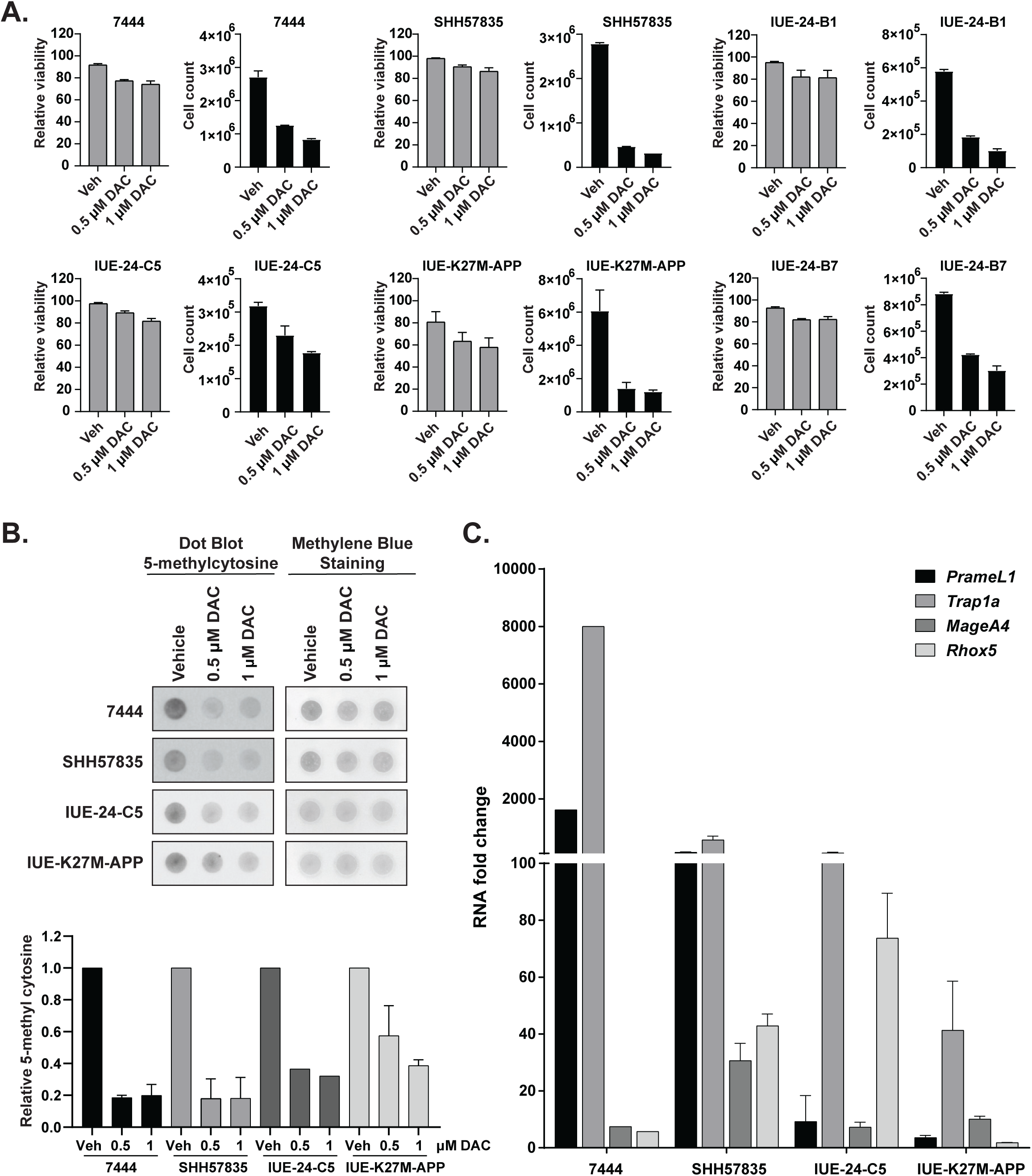
*In vitro* effects of DAC on cell growth, survival, and neoantigen expression in mouse syngeneic cell lines. (A) Cell cytotoxicity was evaluated after treating cells with 0.5 µM, and 1 µM of DAC for 72 hours. (B) 5-methylcytosine-specific dot blot assay was used to evaluate the global hypomethylation. gDNA was isolated from different syngeneic cell lines treated with DAC at two concentrations (0.5 µM and 1 µM) for 72 hours. (Upper panel) 100 ng of gDNA were loaded per dot and methylene blue staining was used as a loading control. (Lower panel) Density of dot blots were measured and plotted as bar graphs. (C) Quantitative PCR evaluation of *PrameL1, Trap1a, MageA4, and Rhox5* neoantigen gene expression in multiple murine syngeneic lines following 3-day 0.5 µM DAC treatment.

### PD-1 inhibition enhances the anti-tumor activity of DAC

Based on the similar effects on growth and neoantigen expression seen after DAC treatment in both PBT and mouse syngeneic lines *in vitro*, we conducted a pilot proof of principle study to determine if DAC could exhibit inhibitory growth effects *in vivo*. IUE-24-C5 DIPG cells were engrafted in the flanks of C57BL/6J (syngeneic line n=4 flanks/group). DAC-treated mice showed reduced tumor volume compared to vehicle mice at 3 weeks post treatment (red vertical hatched line, P=0.0087) **(Supplementary Figure 8A)**. Since previous studies have shown that DAC treatment can increase the efficacy of checkpoint blockade^24^, we further tested the effect of DAC in combination with a murine anti-PD-1 analog of nivolumab, 4H2. At 3 weeks post treatment 4H2 alone did not affect tumor growth (P=0.3817); however, 4H2 in combination with DAC exhibited further tumor suppression than treatment with DAC at this time point (P= 0.0112).

We also evaluated CTA expression and found DAC and DAC+4H2 treated flank tumors showed increased neoantigen and *Irf7* expression (IFN type I induction) **(Supplementary Figure 8B)**. Immunohistochemistry evaluation revealed an increase of CD8^+^ T cells in tumor specimens compared to vehicle after treatment with DAC alone, with the greatest T cell recruitment after the combination of DAC and 4H2 (P= 0.0456 and P= 0.0006, respectively), however, we did not see a statistically significant change in F4/80+ murine macrophages in the flank tumors following any treatment (P= 0.9058 Veh vs. DAC and 0.9510 Veh vs. DAC+4H2) **(Supplementary Figure 8C and D)**. Collectively, the above data confirmed that decitabine induces tumor expression of neoantigens *in vivo* and has anti-tumor activity, and this activity is further enhanced by the addition of an anti-PD-1 antibody.

### DAC treatment induces CTA expression in orthotopic models

Based on the success of our flank model, we next tested the efficacy of DAC with or without 4H2 in a variety of mouse syngeneic orthotopic brain tumor models. Since DAC has been reported to cross the blood-brain barrier, we first wanted to verify that DAC could indeed reach tumors implanted in the brain. Mice were dosed for at least 2 weeks with DAC, then brain tumor tissue was collected, and global DNA methylation activity was quantitated using methylcytosine antibody-based dot blot assay **(Figure 4A, Supplementary Figure 9)**. Compared to vehicle, DAC, and DAC+4H2 treated mice showed a reduction in global methylation of DNA after treatments in all three disease models tested, suggesting that DAC was able to induce epigenetic modulation in multiple locations within the brain at the dose used (0.5 mg/kg). Moreover, we evaluated CTA and *Irf7* expression. DAC and DAC+4H2 treated IUE-24-C5 tumor bearing mice showed induced expression of *MageA4, Trap1a,* and *Rhox5*. DAC alone and its combination with 4H2 also activated the expression of *Irf7*, suggesting the induction of the IFN type I pathway **(Figure 4B)**.

**Figure 4.**
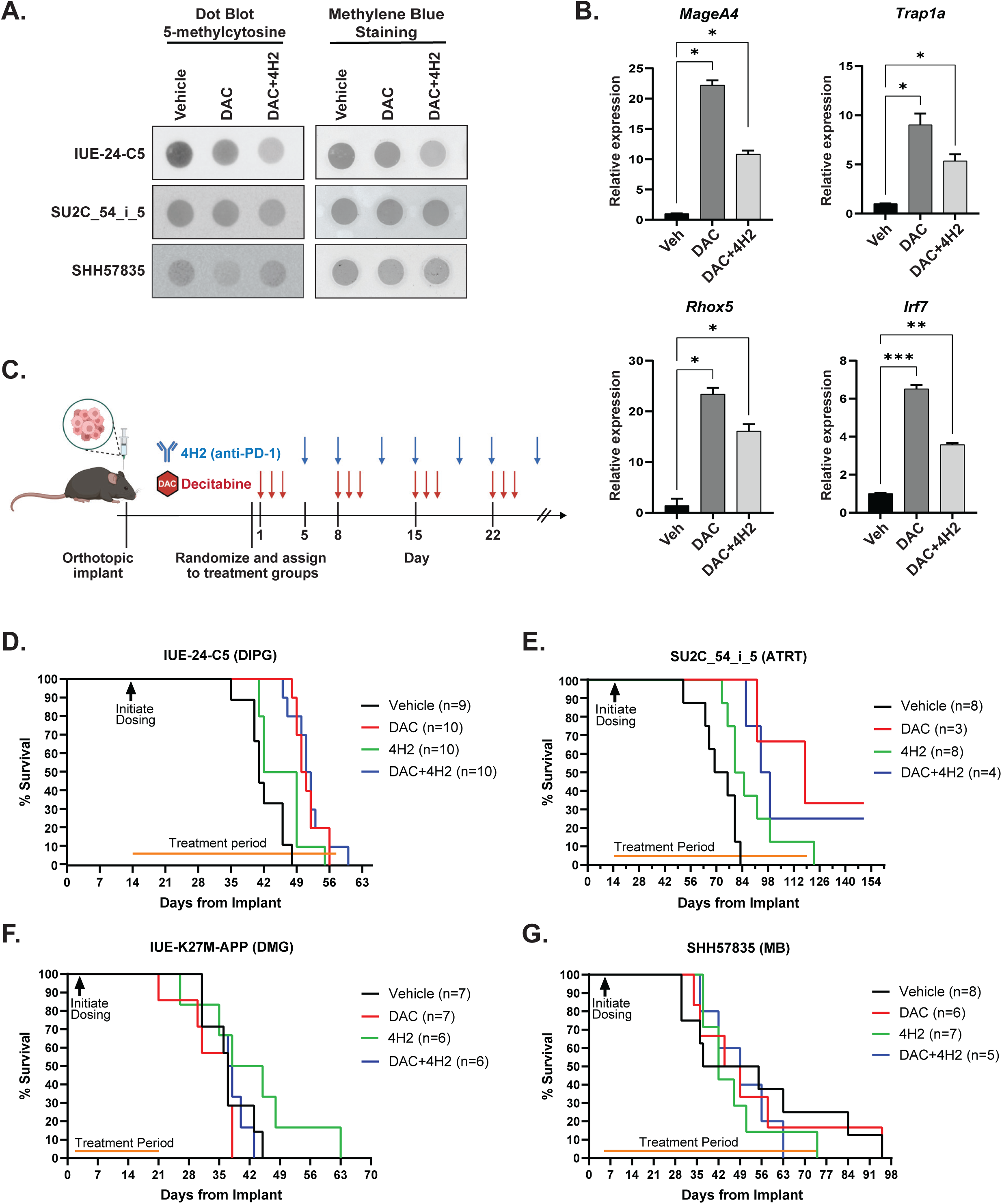
Variable responses to DAC, DAC+4H2, or 4H2 treatment in orthotopic syngeneic models. (A) 5-methylcytosine-specific dot blot assay was used to evaluate activity of DAC in mouse brain tumor tissue. gDNA was isolated from brain tumors established from different models and treated with DAC or a combination of DAC and 4H2. 100 ng of gDNA were loaded per dot and methylene blue staining was used as a loading control. (B) Evaluation of neoantigen gene expression and inflammation using quantitative PCR in tumor tissue harvested from IUE-24-C5 syngeneic mice treated with either vehicle (Veh), DAC, or DAC+4H2. (C) Schematic representation of treatment strategy of DAC and 4H2 in syngeneic models. Red arrows indicate DAC IP dosing (0.5 mg/kg) and blue arrows indicated anti-PD-L1 (4H2) IP dosing (200 μg). Figure created with BioRender.com. Kaplan-Meier curves representing survival data for (D) DIPG model IUE-24-C5, (E) ATRT model SU2C_54_i_5, (F) DMG model IUE-K27M-APP, and (G) MB model SHH57835.

### DAC and its combination with 4H2 improved survival in DIPG and ATRT models

With the knowledge that DAC can reach orthotopic tumors, we initiated a treatment regimen to determine if DAC, DAC+4H2, or 4H2 alone could improve survival in mouse brain tumor models **(Figure 4C)**. In the IUE-24-C5 syngeneic model, we injected 100,000 cells into the pons using stereotactic coordinates. Mice were randomized into 4 groups (Vehicle n=9, DAC n=10, 4H2 n=10, DAC+4H2 N=9) based on tumor establishment using bioluminescence imaging data. Tumor measurements show that during the initial phase of tumor growth, tumors treated with DAC alone or DAC+4H2 grew more slowly compared to vehicle-treated mice **(Supplementary Figure 10)**. However, tumors progressed at later stages and mice eventually succumbed to disease in all treatment arms **(Figure 4D)**. As measured by Kaplan-Meier survival analysis, there was a significant overall survival (OS) benefit in the DAC and DAC+4H2 combination group (OS in days, Veh=41, DAC=50.5, 4H2=45.5, DAC+4H2=51.5) (p<0.0001 and 0.0170 respectively compared to vehicle for DAC alone and DAC+4H2 combination) **(Figure 4D).** Overall, DAC inhibits tumor growth in the IUE-24-C5 orthotopic model but the combination of DAC+4H2 did not show superiority compared to DAC monotherapy. Unfortunately, we could not replicate the study on histone 3.1 and 3.3 mutant mouse model DMG tumors due to the slow tumor latency of these models. For the ATRT model (SU2C_54_i_5), mice were randomized after confirmation of tumor establishment on MRI **(Supplementary Figure 11)**. Kaplan-Meier survival analysis showed a significant OS benefit with DAC, DAC+4H2, and 4H2 alone (OS in days, Veh=72.5, DAC=118, DAC+4H2=96.5, 4H2=82.5) (p=0.0079, 0.0027, 0.0176 respectively to vehicle for DAC alone, DAC+4H2 combination, 4H2 alone) **(Figure 4E)**.

We also tested a DMG model (IUE-K27M-APP) **(Figure 4F)** and two different medulloblastoma models (SHH57835, 7444) **(Figure 4G, Supplementary Figure 12)**. In contrast to the DIPG and ATRT models, these models failed to show significant survival benefit regardless of treatment.

### DAC+4H2 treatment can remodel the immune tumor microenvironment

Variable responses to DAC+4H2 treatment across models prompted us to employ CyTOF-based immune cell analysis to evaluate the cell types present in the TME at baseline and after treatment **(Figure 5, Supplementary Figure 13)**. Baseline analysis of models have 32 clusters of immune populations distributed among 8 different cell types. Clusters among different populations were distributed as 10 in macrophages, 4 in neutrophils, 5 in T cells, 5 in MDSCs, 2 in DCs, 2 in monocytes, 12 in NK cells, and 2 were unassigned **(Figure 5A and B)**. Individual t-SNE plots showed significant variability at baseline between the tumor models **(Figure 5C)**. HGG and ATRT models showed higher T cell infiltration compared to other models. MDSCs populations were significantly high in IUE-24-C5 and ATRT models suggesting the crucial role in immune suppression. In addition, IUE-24-C5 and ATRT models had a significant fraction of macrophages with CCR2+ supporting the immunosuppressive microenvironment shown previously.^52^ This variability in cell type clusters changed even more upon DAC+4H2 treatment among all models **(Figure 6)**. Analysis of leukocyte markers showed changes in the immune composition of the tumors upon treatment **(Figure 6A)**. Specifically, in the DIPG IUE-24-C5 model, significant differences were observed in the macrophage, T cell, and MDSC populations upon treatment **(Figure 6B)**. Macrophage and MDSC markers decreased, while CD4+ T cells significantly increased. This observation supports the survival benefit observed in the IUE-24-C5 model **(Figure 4D)**. Analysis within each cluster showed even more changes **(Figure 6C)**. MDSC clusters 5, 24, 29, along with macrophage cluster 22 and neutrophil cluster 18 showed significant decrease. In contrast, cluster 16 (CD4+ T cells PD1+) was significantly increased upon combination treatment of the IUE-24-C5 tumor bearing mice.

**Figure 5.**
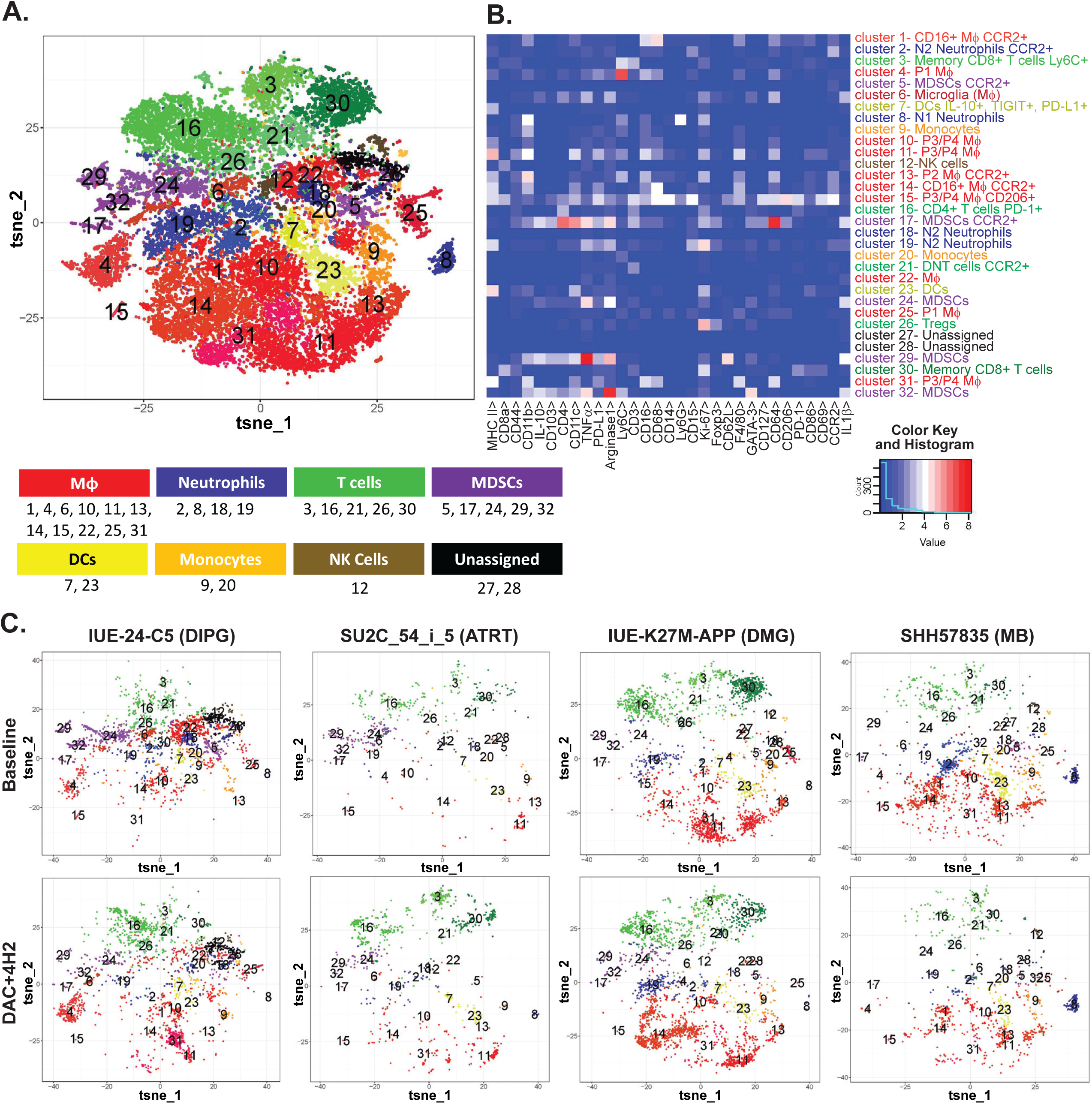
Immune composition of syngeneic TME at baseline vs. DAC+4H2 combination treatment. The infiltration of immune cells and effect of combination treatment was analyzed using CyTOF. (A) Phenograph clustering of all samples. A t-SNE plot representing all models at baseline with different cell populations as clusters indicated by color. (B) Heat map showing expression of specific immune markers in the various cluster groups. (C) t-SNE plots for each individual tumor model at both baseline (upper) and following DAC+4H2 (lower) treatment (n=3 tumors per condition, per model).

Similar differences were observed in the ATRT model, where the MDSC clusters 24 and 29 were decreased, while an increase in neutrophils was seen, suggesting an inflammatory response **(Figure 6B and D)**. In contrast, IUE-K27M-APP and SHH57835 models had comparably low MDSC and CCR2+ populations at baseline and further DAC+4H2 treatment did not change the composition significantly suggesting these models have independent mechanisms to repress the immune landscape of these tumors. Of note, small but significant changes were seen in clusters 9, 20, 23, and 31 from DMG tumors **(Figure 6E)** and clusters 17 and 24 from SHH57835 tumors **(Figure 6F)**. These small changes possibly reflect the lack of efficacy seen in both the DMG (IUE-K27M-APP) and the medulloblastoma (SHH57835) models.

**Figure 6.**
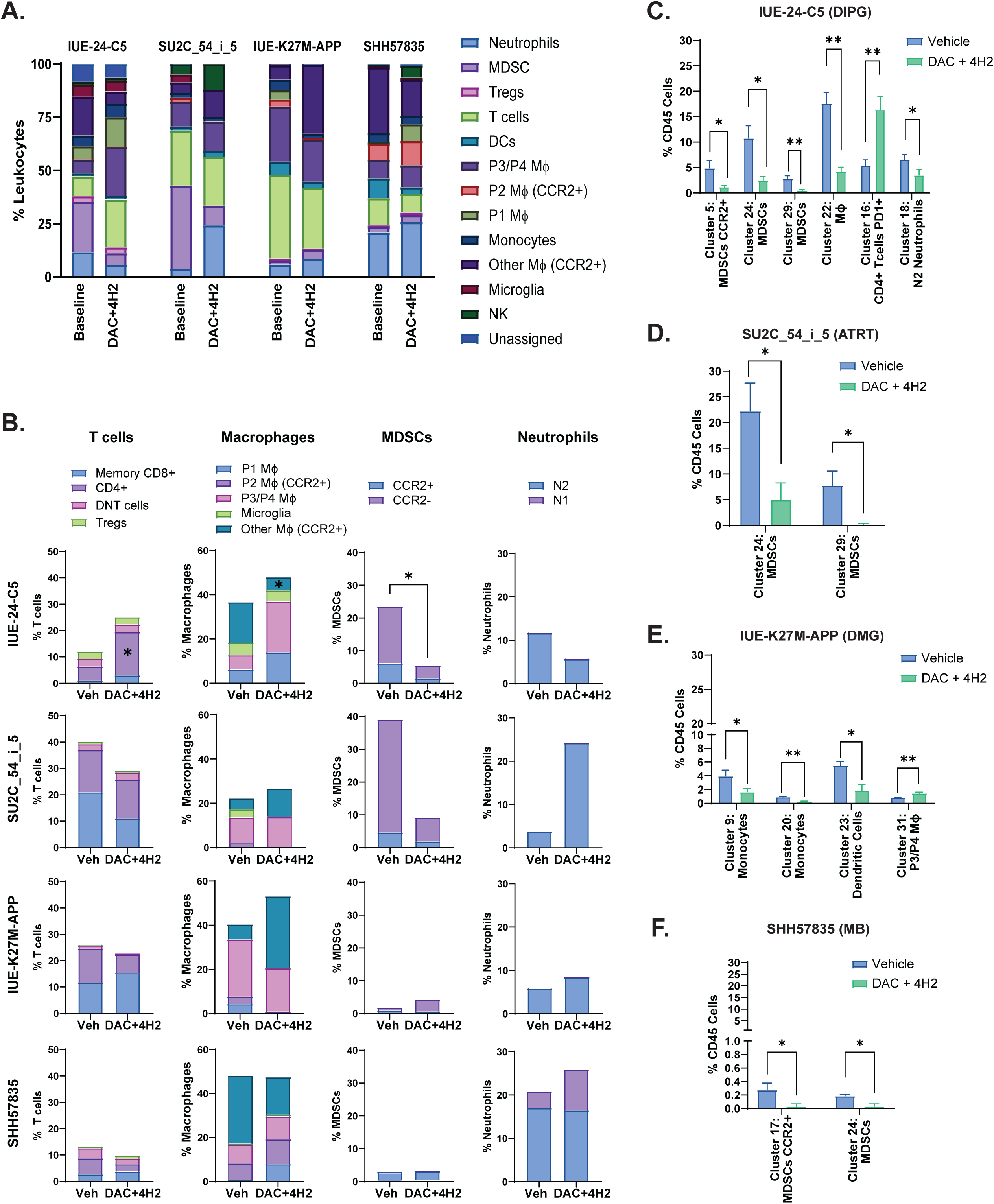
Cluster analysis of syngeneic tumors at baseline vs. DAC+4H2 treatment. (A) Relative proportions of leukocyte populations in each tumor type at baseline and following DAC+4H2 treatment. (B) Relative percentage of T cells, macrophages, MDSCs, and neutrophils in each tumor model from vehicle treated mice and those treated with DAC+4H2. Significant changes in individual cell clusters for (C) IUE-24-C5 (D) SU2C-54_i_5 (E) IUE-K27APP and (F) SHH57835 models. *P* = 0.001 - 0.01 **, 0.01 - 0.05 *

## DISCUSSION

The unique tumor microenvironment (TME) present in PBTs features a low lymphocytic infiltrate and an immunologically inert landscape that is associated with a low mutational burden. This unique TME, coupled with poor BBB-penetration of therapeutic antibodies, helps explain the historical sub-optimal responses observed following ICI therapy in PBTs. Indeed, there have been durable therapeutic outcomes in a subset of rare pediatric tumors with high mutational burden that demonstrate remarkable response to ICI therapy.^12,53,54^ Therefore, there is a significant opportunity for immune priming agents to aid and address this discrepancy in order to provide increased efficacy of ICI therapy in PBTs.

Our findings suggests that utilization of DAC, as an immune priming agent, may provide benefit in DIPG and ATRT syngeneic murine models. Additionally, we noted a prolonged survival benefit and TME differences in the macrophage, MDSC, and T-cell populations upon treatment compared to control in the DIPG and ATRT models respectively. Interestingly, both the DIPG and ATRT models had a high baseline MDSC population prior to treatment with DAC and 4H2 that was markedly reduced with treatment. This was in stark contrast to the non-responding DMG and MB models which did not contain as high MDSC populations, indicating potential differences between tumor subtypes as an important consideration for immune priming agents. The outcomes observed in both DIPG and ATRT murine models suggest that combining an immune-priming agent with ICI could represent a viable therapeutic approach to improve patient outcomes and improve the efficacy of ICI agents.

MDSCs, a heterogenous group of mature and immature monocytic and granulocytic lineage cells released from the bone marrow, have long been associated with tumor progression and immune evasion due to their ability to suppress CD4^+^ and CD8^+^ T-cell proliferation.^55^ Similar to observations in murine ovarian cancer models, utilizing an agent like DAC which can modulate MDSC activity, may reduce lymphocytic suppression in the microenvironment produced by MDSCs in PBTs.^56,57^ Additionally, solid tumors are often engulfed by macrophage populations, of which a large proportion is MDSCs. CCR2+ monocytic cells inhibit the T cell infiltration in solid tumors and our data in IUE-24-C5 and ATRT model shows a similar observation.^52^ This suggests that within higher-grade tumors, where MDSC expression is likely increased, reversing the immune suppressive state driven MDSCs would be essential for the efficacy of ICI therapy.

Conversely, another approach involves promoting a greater immunogenic TME. Proposed strategies include utilizing agents that promote endogenous retroviral expression and increase the Type I/II interferon response to drive PD-1 expression and increase tumor-antigen presentation to the host immune cell.^20^ While decitabine has known endogenous retroviral expression activity, our findings suggest that its anti-MDSC effects produced greater ICI affinity than its pro-immunogenic effects, suggesting that it was important to promote a pro-lymphocytic proliferation TME initially rather than enhance tumor immunogenicity.

While our findings suggest a promising initial step in enhancing the efficacy of ICI therapy in PBT, several challenges and limitations remain. The translation from a syngeneic murine model to clinical trials may not reflect the complexity of the biology and TME that is present in our patient cohort. For instance, it is likely that there will be increased MDSC populations across the board in PBTs due to the chronicity of their tumor growth compared to the shorter growth time present in murine models. Therefore, if targeting MDSC populations improves ICI efficacy, this suggests that there could be an opportunity to provide varying patient benefit across the board through immune priming in PBTs. However, while targeting MDSC populations does show promise as a strategy to generate a greater immunogenic response in PBTs, the role of other immunosuppressive mechanisms that exist within the TME should not be overlooked. Tumors have been shown to employ a vast array of evasion and suppression mechanisms including regulatory T-cells, tumor-associated macrophages and immunosuppressive cytokines which may provide additional therapeutic targets for immune priming and improvement of ICI therapy. Therefore, further investigation is required to assess alternate immunological targets that may render non-responding tumors, such as the DMG and MB models in our study, responsive.

Second, blood-brain barrier (BBB) penetration of ICI molecules remains a consideration. Mouse cancer models including ovarian^56^, colon^24,58^, pancreatic^59^, and melanoma^60^ have shown remarkable preclinical efficacy when treated with DNMTi and ICIs. However, these models don’t have the added BBB complexity to contend with and thus the effective concentrations of ICIs reaching their target are likely higher. Current preclinical and clinical studies are actively addressing this issue to increase the delivery of ICIs to brain tumor cells. Techniques like convection-enhanced or ultrasound-mediated delivery, laser interstitial thermotherapy, along with optimal timing and sequencing strategies of surgery and/or radiation and administration of ICIs hold promise to lower this barrier.^61^

Lastly, the potential adverse effects of combination immune-priming agents and ICI therapy requires further investigation. While DAC can enhance anti-tumor immune responses by modulating the TME, its systemic effects are capable of producing a myelosuppressive state and reduce hematopoiesis.^62^ Alongside the well-established immunological inflammatory side effects and pseudo-progression that are associated with ICI therapy, it remains unknown what the adverse effect profile will be in patients who have been administered both agents. Balancing the therapeutic benefits against the adverse effects is essential for optimizing the therapeutic regimen and ensuring patient safety. Moreover, DNMTi have dual activity, hypomethylation of DNA and cytotoxic activity via DNA damage.^63^ This could inhibit the proliferation of immune cells. Therefore, the selection of optimum dosing and timing is important for sustained hypomethylation to achieve the intended immunomodulatory effect.

In conclusion, we show that through analysis of various pediatric tumor gene expression data sets that there is heterogenous expression of various immune associated markers. Specifically, the heterogeneity of MHC I and PD-L1 can also be observed in various patient-derived cell lines. We also demonstrate that low-dose DAC treatment can decrease growth and increase MHC I and neoantigen expression in human tumor cell lines *in vitro*. Furthermore, we found that these responses to low-dose DAC treatment translate to mouse syngeneic cell lines, representing various PBTs, which allow us to preclinically test various treatments in animals with intact immune systems. Although DAC induced changes in DNA methylation and neoantigen expression in orthotopic allografts, only the DIPG and ATRT models showed improved survival. This differential response may be partially explained by changes seen in the immune cell composition of the TME, particularly MDSCs, thus providing some insight into possible ways to stratify patients for future clinical trials.

## REQUIRED STATEMENTS

### Ethics

All mouse experiments were conducted in accordance with the National Institutes of Health (NIH) *Guide for the Care and Use of Laboratory Animals*, with approval from the Institutional Care and Use Committees (IACUC) at each site.

### Authorship

Conceptualization: M.F., N.J., A.H., J.M.O., R.D.; Experimental design: D.K.M., S.M.M., D.P., E.J.G., A.B., S.S.K., A.F.A., M-Y.B., E.E.C., K.B., C.R., M.Z., M.C.B., M.C., D.L-L., L.K., M.F., N.J., A.H., J.M.O., R.D.; Data acquisition and analysis: D.K.M., S.M.M., D.P., E.J.G., A.B., S.S.K., A.F.A., X.Y., F.Y., M-Y.B., U.B., E.E.C., K.B., F.P., C.R., M.Z., M.C.B., M.C., D.L-L., L.K.; Study supervision: N.J., A.H., J.M.O., R.D.; Writing, editing & review: all authors. All authors have read and agreed to the published version of the manuscript.

### Data Availability

All data generated in this study are available upon request from the corresponding authors.

## Acknowledgements

The authors would like to thank Bristol Myers Squibb for generously providing the anti-PD-1 antibody, corresponding isotype control, and experimental guidance for these studies; Stand Up to Cancer, The Cure Starts Now Foundation, and The Canadian Institute of Health Research for their financial and scientific support; and all the patients and families that inspire us to find better treatments. We also thank Drs. Martine Roussel, Michelle Monje, Esther Hulleman, Tim Phoenix, Manav Pathania, and Paolo Salomoni for kindly providing various human and mouse cell lines, Jie Wang and Morgan Merrill for their technical support, Sara Lawellin and Michelle Deutsch for administrative support.

## SUPPLEMENTAL MATERIALS

### Cell Lines

| **Human Cell Lines** | **Tumor Type** | **Source** |
| --- | --- | --- |
| PBT-14FHTC | DMG, H3.3K27M | Olson<sup>36</sup> |
| PBT-22FHTC | DMG, H3.3K27M | Olson<sup>36</sup> |
| PBT-29FHTC | DMG, H3.3K27M | Olson<sup>36</sup> |
| SU-DIPG-XIII | DMG, H3.3K27M | Monje<sup>37</sup> |
| SU-DIPG-XVII | DMG, H3.3K27M | Monje<sup>37</sup> |
| PBT-27FHTC | DMG, H3.1K27M | Olson<sup>36</sup> |
| SU-DIPG-IV | DMG, H3.1K27M | Monje<sup>37</sup> |
| CCHMC-DIPG-1 | DIPG, H3WT | Drissi<sup>38</sup> |
| PBT-24FHTC | DIPG, H3WT | Olson<sup>36</sup> |
| VUMC-DIPG-10 | DIPG, H3WT | Hulleman |
| Med-114FHTC | Medulloblastoma, Group 3, *MYC* amp | Olson<sup>39</sup> |
| Med-411FHTC | Medulloblastoma, Group 3, *MYC* amp | Olson<sup>39</sup> |
| Med-2112FHTC | Medulloblastoma, Group 3, *MYC* amp | Olson<sup>39</sup> |
| Med-813FHTC | Medulloblastoma, Group 3, *MYC* amp | Olson<sup>39</sup> |
| ATRT-311FHTC | ATRT, SHH | Olson<sup>39</sup> |
| ATRT-310FHTC | ATRT, SHH | Olson<sup>39</sup> |
| CHLA-04 | ATRT, SHH | ATCC |
| BT-12 | ATRT, MYC | Children's Oncology Group<sup>40</sup> |
| BT-16 | ATRT, MYC | CHOP<sup>41</sup> |
| CHLA-06 | ATRT, MYC | ATCC |
| PBT-05FHTC | HGG, *MYC-N* amp | Olson<sup>39</sup> |
| GBM-110FHTC | HGG | Olson<sup>39</sup> |
| GBM-511FHTC | HGG | Olson<sup>39</sup> |
| SU-pcGBM2 | HGG | Monje<sup>37</sup> |
| EPD-210FHTC | Ependymoma | Olson<sup>39</sup> |

| Murine Cell Lines | Tumor Type | Source |
| --- | --- | --- |
| IUE-24-C5 | *Pdgfra*<sup>D842V</sup>, dominant neg<br>*Trp53*, H3 WT | Phoenix<sup>42</sup> |
| IUE-K27M-APP | H3.3K27M, *Trp53-Atrx*-KD,<br>PDGFRA-overexpression | Salomoni<sup>34</sup> |
| IUE-24-B1 | *Pdgfra*<sup>D842V</sup>, dominant neg<br>*Trp53*, H3.3K27M | Phoenix, Hulleman<sup>32</sup> |
| IUE-24-B7 | *Pdgfra*<sup>D842V</sup>, dominant neg<br>*Trp53*, H3.3K27M | Phoenix, Hulleman<sup>32</sup> |
| SHH57835 | *Ptch1*<sup>+/-</sup>, *Trp53* KO | Roussel<sup>33</sup> |
| 7444 | *c-Myc* CRISPR, *Trp53* loss,<br>*Cdkn2c* loss | Roussel<sup>35</sup> |

### qPCR Primers

**qPCR Primers:**
| Gene Target | Taqman Assay |
| --- | --- |
| *CTAG1B (NY-ESO-1)* | Hs00265824_m1 |
| *MAGEA1* | Hs00607097_m1 |
| *ACTB* | Hs99999903_m1 |
| *PrameL1* | Mm00473193_m1 |
| *Trap1a* | Mm00495785_m1 |
| *MageA4* | Mm00522322_s1 |
| *Rhox5* | Mm00476718_m1 |
| *Irf7* | Mm00516788_m1 |
| *GusB* | Mm01197698_m1 |

### Antibodies

**Antibodies:**
| **Target** | **Source** | **Application** |
| --- | --- | --- |
| 5-methylcytosine | Sigma Aldrich (MABE146) | Dot Blot |
| PE Mouse IgG1, κ Isotype control | BioLegend (400114) | Flow |
| PE Mouse IgG2a, κ Isotype control | BioLegend (400214) | Flow |
| APC Mouse IgG2a, κ Isotype control | BioLegend (400220) | Flow |
| APC Rat IgG2a, κ Isotype control | BioLegend (400512) | Flow |
| PE anti-human HLA-A,B,C | BioLegend (311406) | Flow |
| APC anti-human HLA-A,B,C | BioLegend (311410) | Flow |
| PE anti-human CD274 (B7-H1, PD-L1) | BioLegend (329706) | Flow |
| APC anti-human CD274 (B7-H1, PD-L1) | BioLegend (329708) | Flow |
| APC anti-mouse H-2K<sup>b</sup>/H-2D<sup>b</sup> | BioLegend (114614) | Flow |
| PE anti-mouse CD274 (B7-H1, PD-L1) | BioLegend (124308) | Flow |
| CD3 | Roche (790-4341),<br>Abcam (ab11089, ab16669) | IHC |
| CD4 | Abcam (ab183685) | IHC |
| CD8 | Cell Signaling (98941),<br>Abcam (ab209775) | IHC |
| F4/80 | Abcam (ab6640) | IHC |
| PD-1 | Abcam (ab214421) | IHC |
| FoxP3 | ThermoFisher (14-5773-82) | IHC |

### CyTOF Antibodies

**CyTOF Antibodies:**
| **Antibody** | **Metal** | **Cell Types/Function** |
| --- | --- | --- |
| CD45 | 89Y | Leukocytes |
| MHCII (IAIE) | 113Cd | APC |
| CD8a | 116Cd | Cytotoxic T cells |
| CD44 | 141Pr | Memory |
| CD11b | 142Nd | Myeloid |
| IL-10 | 143Nd | Cytokine (anti) |
| CD103 | 144Nd | Tissue residency |
| CD4 | 145Nd | Helper T cells |
| CD11c | 146Nd | Dendritic Cells |
| CD223/LAG3 | 147Sm | Check point inhibitor |
| TNFα | 148Nd | Cytokine (pro) |
| PD-L1 | 149Sm | Check point inhibitor |
| Arginase 1 | 150Nd | Anti-inflammatory |
| Ly6C | 151Eu | Myeloid/monocytes |
| CD3 | 152Sm | T cells |
| CD16 | 153Eu | IgG receptor myeloid |
| CD284/TLR4 | 154Sm | Innate signaling |
| CD68 | 155Gd | Monocytes/macrophage |
| CD14 | 156Gd | Monocytes |
| Ly6G | 158Gd | Neutrophils |
| CD15 | 159Tb | Neutrophils |
| Ki-67 | 161Dy | Proliferation |
| Foxp3 | 162Dy | Tregs |
| TIGIT | 163Dy | Check point inhibitor |
| CD62L | 164Dy | Naïve |
| F4/80 | 165Ho | Macrophage |
| GATA-3 | 166Er | Transcription factor |
| CD127 | <sup>167</sup>Er | IL7 R, ILCs T cells |
| CD64 | <sup>168</sup>Er | Macrophage |
| CD206 | <sup>169</sup>Tm | Anti-inflammatory Macrophage |
| RoRγt | <sup>170</sup>Er | Transcription factor |
| PD-1 | <sup>171</sup>Yb | Check point inhibitor |
| CD86 | <sup>172</sup>Yb | Pro-inflammatory Macrophage |
| CD69 | <sup>173</sup>Yb | Activation/residency |
| CCR2 | <sup>174</sup>Yb | Recent infiltration |
| Tbet | <sup>175</sup>Lu | Transcription factor |
| IL1β | <sup>176</sup>Yb | Cytokine (pro) |

## SUPPLEMENTAL FIGURES

**S1.**
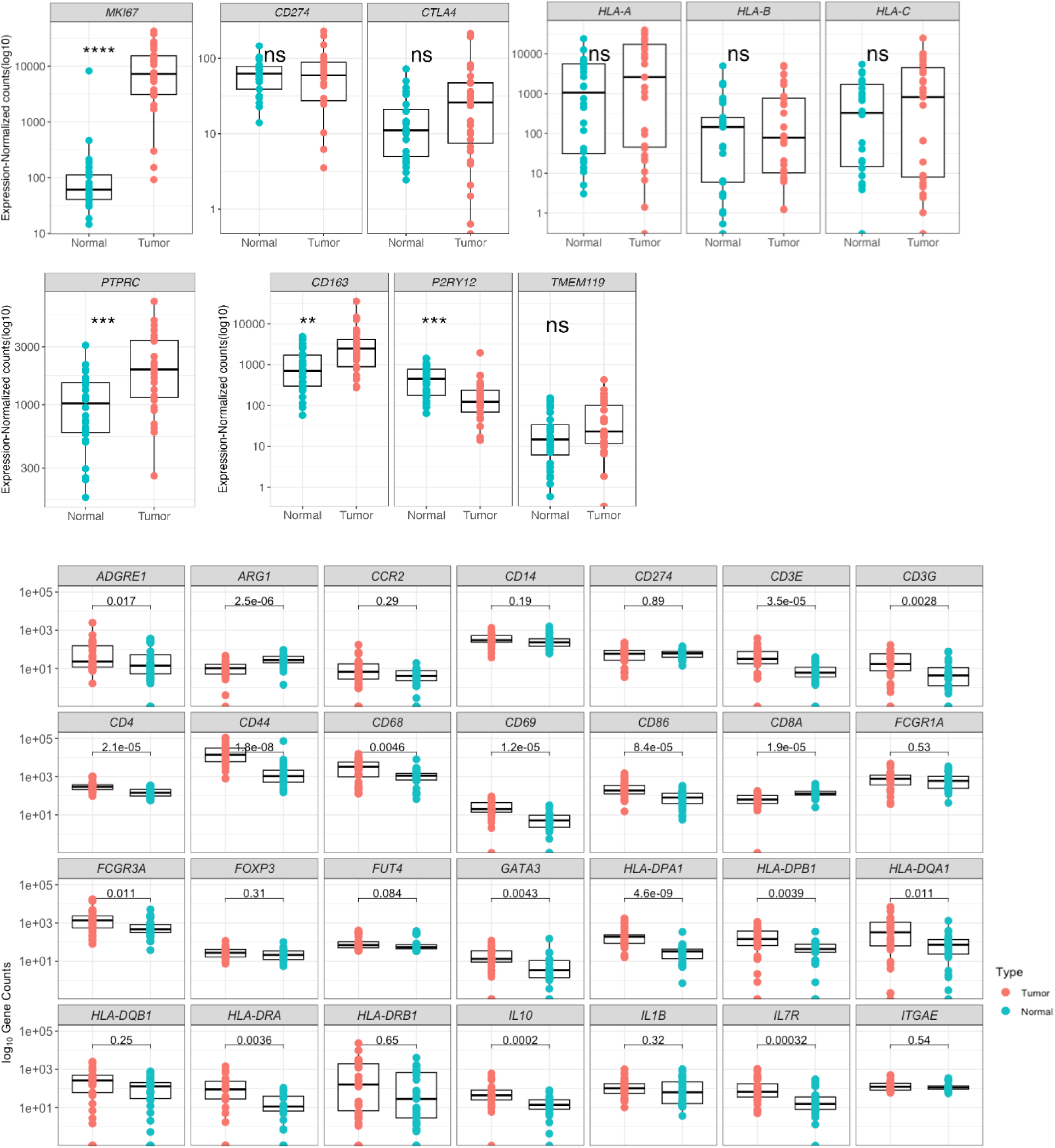
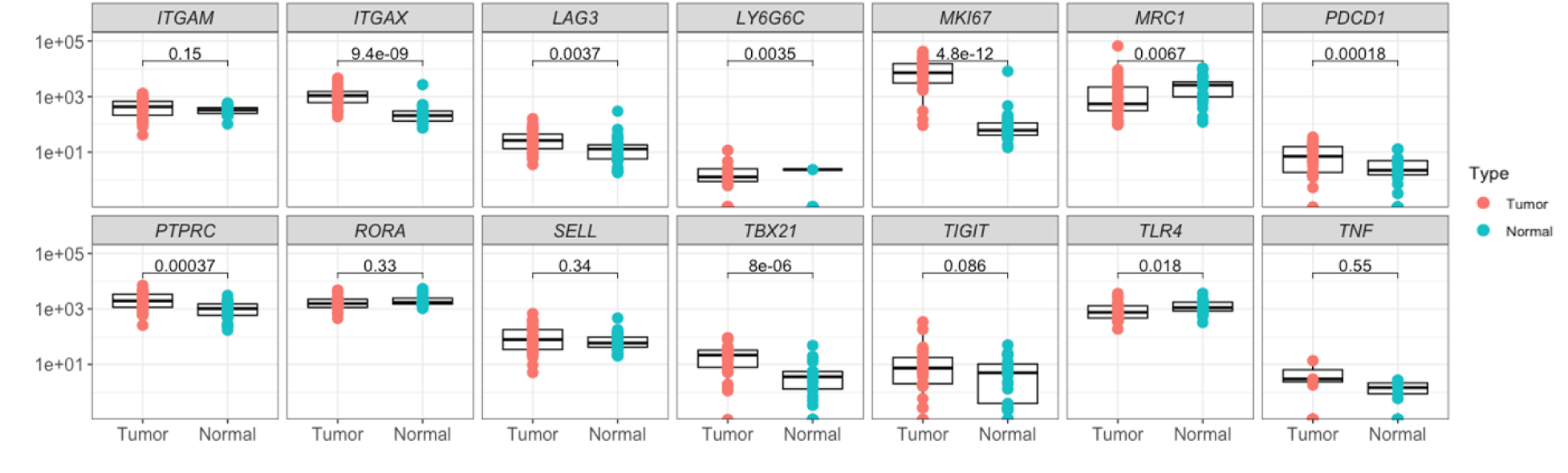
Select immune marker gene expression in DIPG/DMG tumor samples compared to normal brain. Stars represent: * *P* ≤ 0.05, ** *P* ≤ 0.01, *** *P* ≤ 0.001, **** *P* ≤ 0.0001. ns = *P* > 0.05.

**S2.**
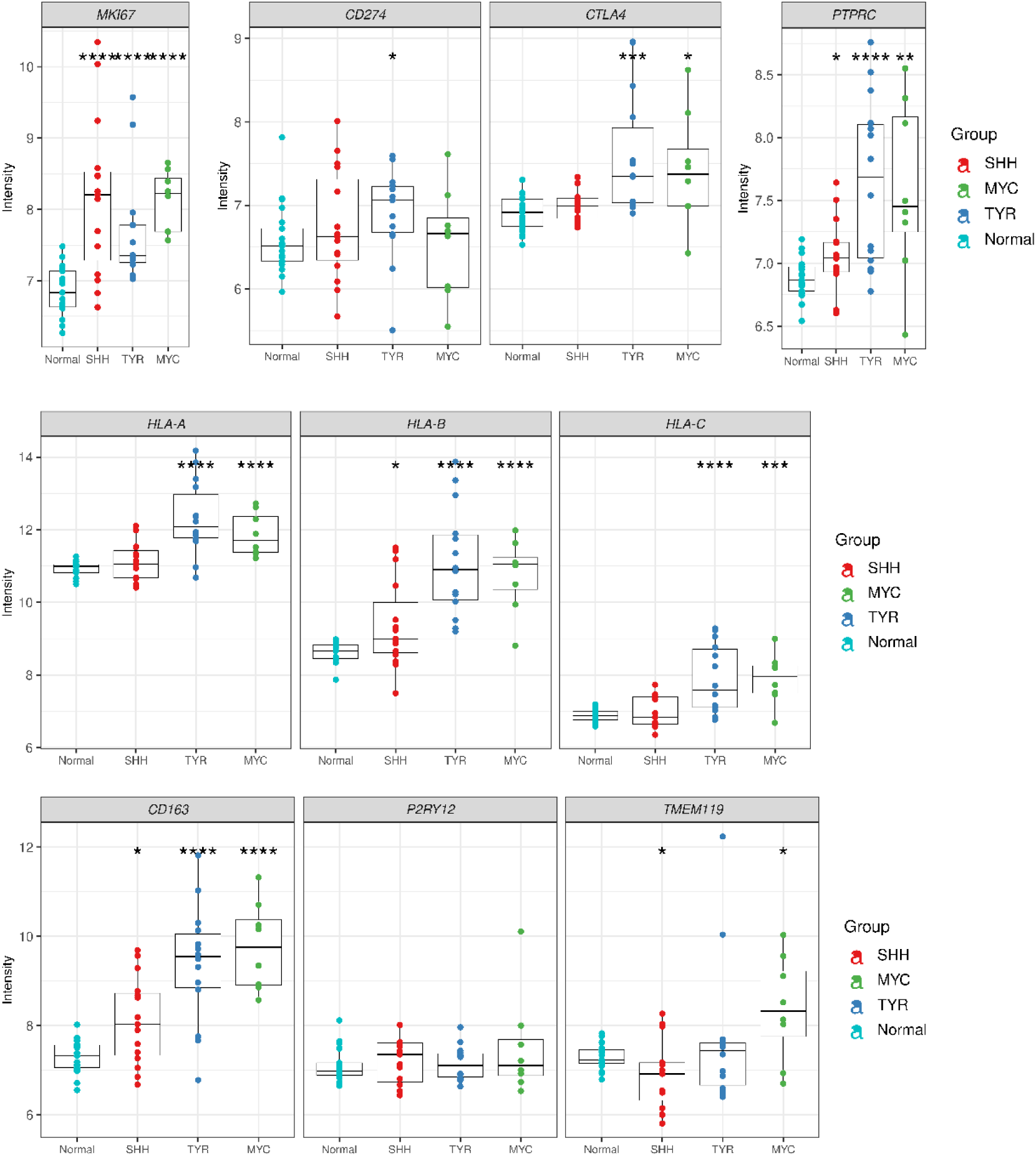
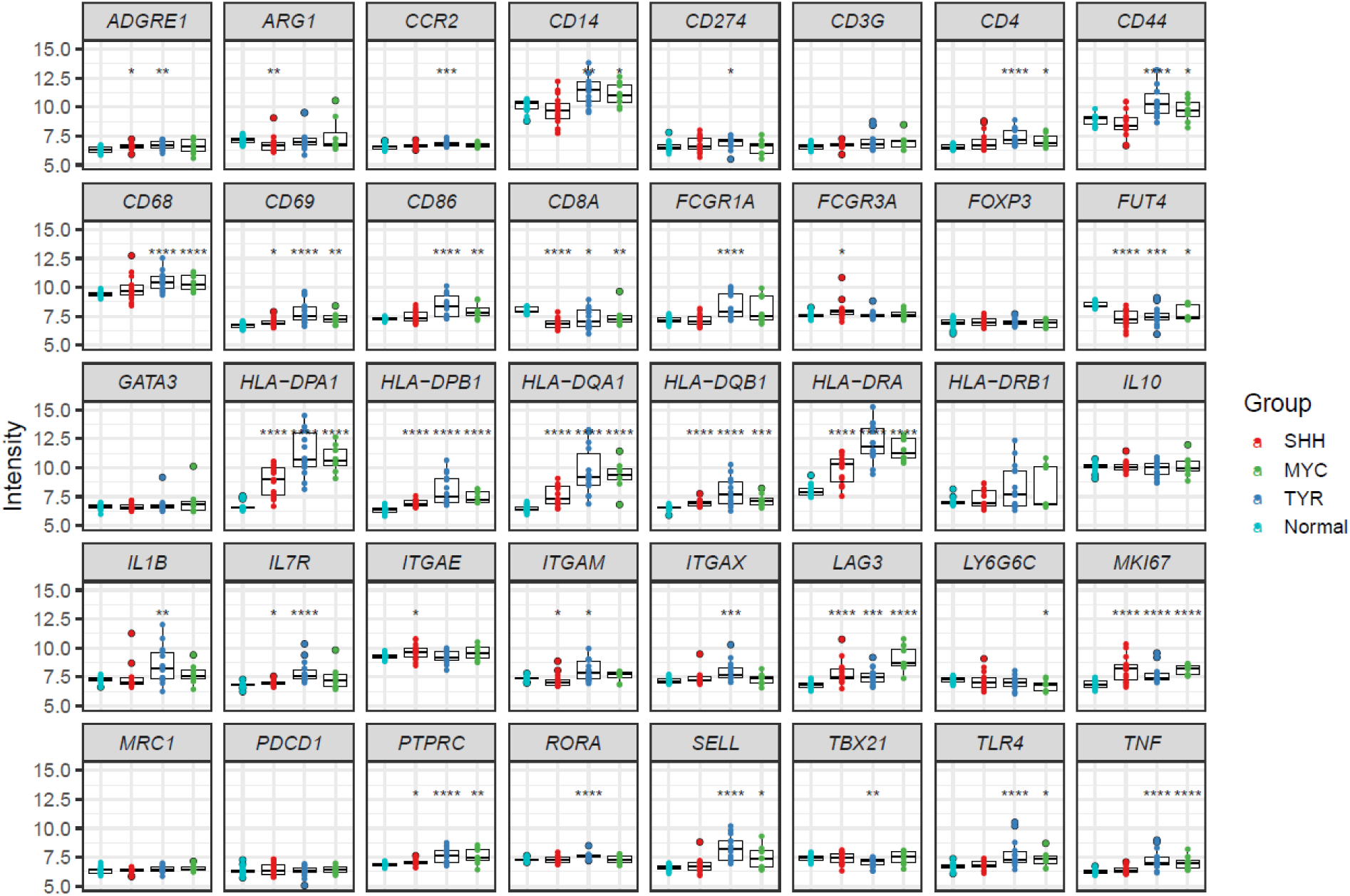
Select immune marker gene expression in ATRT tumor samples compared to normal brain. Stars represent: * *P* ≤ 0.05, ** *P* ≤ 0.01, *** *P* ≤ 0.001, **** *P* ≤ 0.0001.

**S3.**
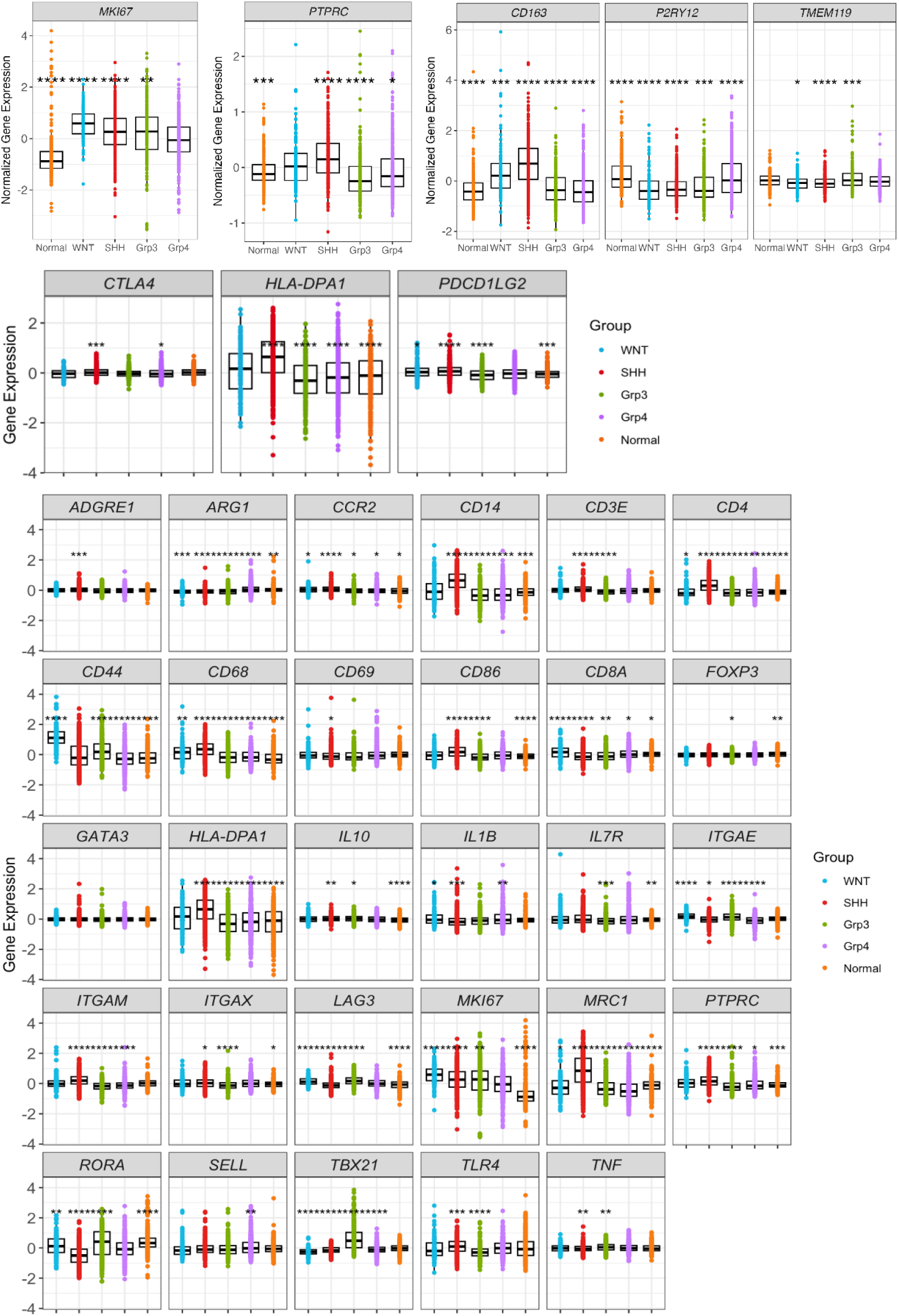
Select immune marker gene expression in medulloblastoma tumor samples compared to normal brain. From Swartling data set (Weishaupt H, Johansson P, Sundstrom A, et al. Batch-normalization of cerebellar and medulloblastoma gene expression datasets utilizing empirically defined negative control genes. *Bioinformatics.* 2019; 35(18):3357-3364). Stars represent: * *P* ≤ 0.05, ** *P* ≤ 0.01, *** *P* ≤ 0.001, **** *P* ≤ 0.0001.

**S4.**
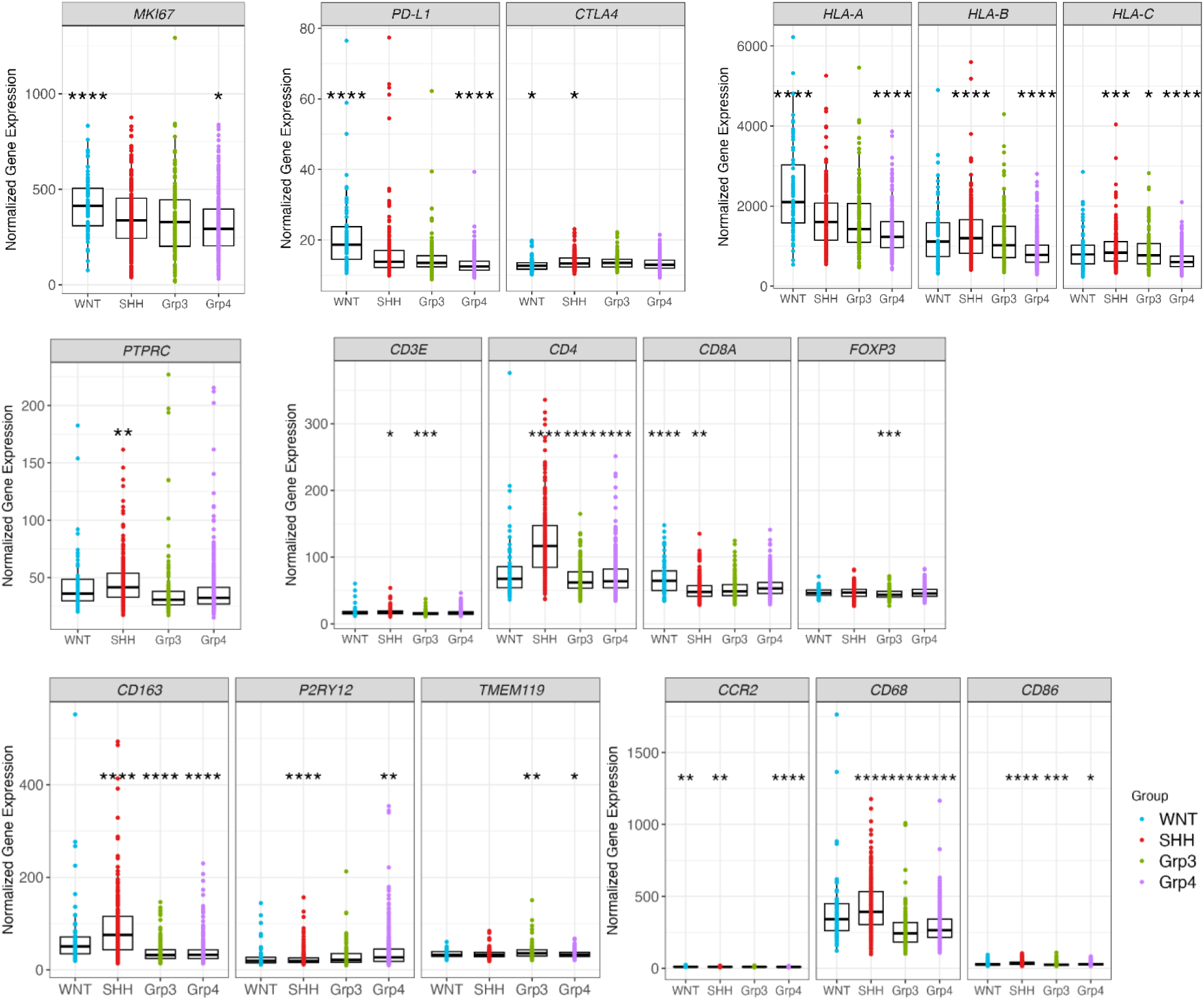
Select immune marker gene expression in medulloblastoma tumor samples. From Cavalli data set (Cavalli FMG, Remke M, Rampasek L, et al. Intertumoral Heterogeneity within Medulloblastoma Subgroups. *Cancer Cell.* 2017; 31(6):737-754 e736). Data set includes 763 primary samples consisting of 70 WNT, 223 SHH, 144 group 3, and 326 group 4 samples. Stars represent: * *P* ≤ 0.05, ** *P* ≤ 0.01, *** *P* ≤ 0.001, **** *P* ≤ 0.0001.

**S5.**
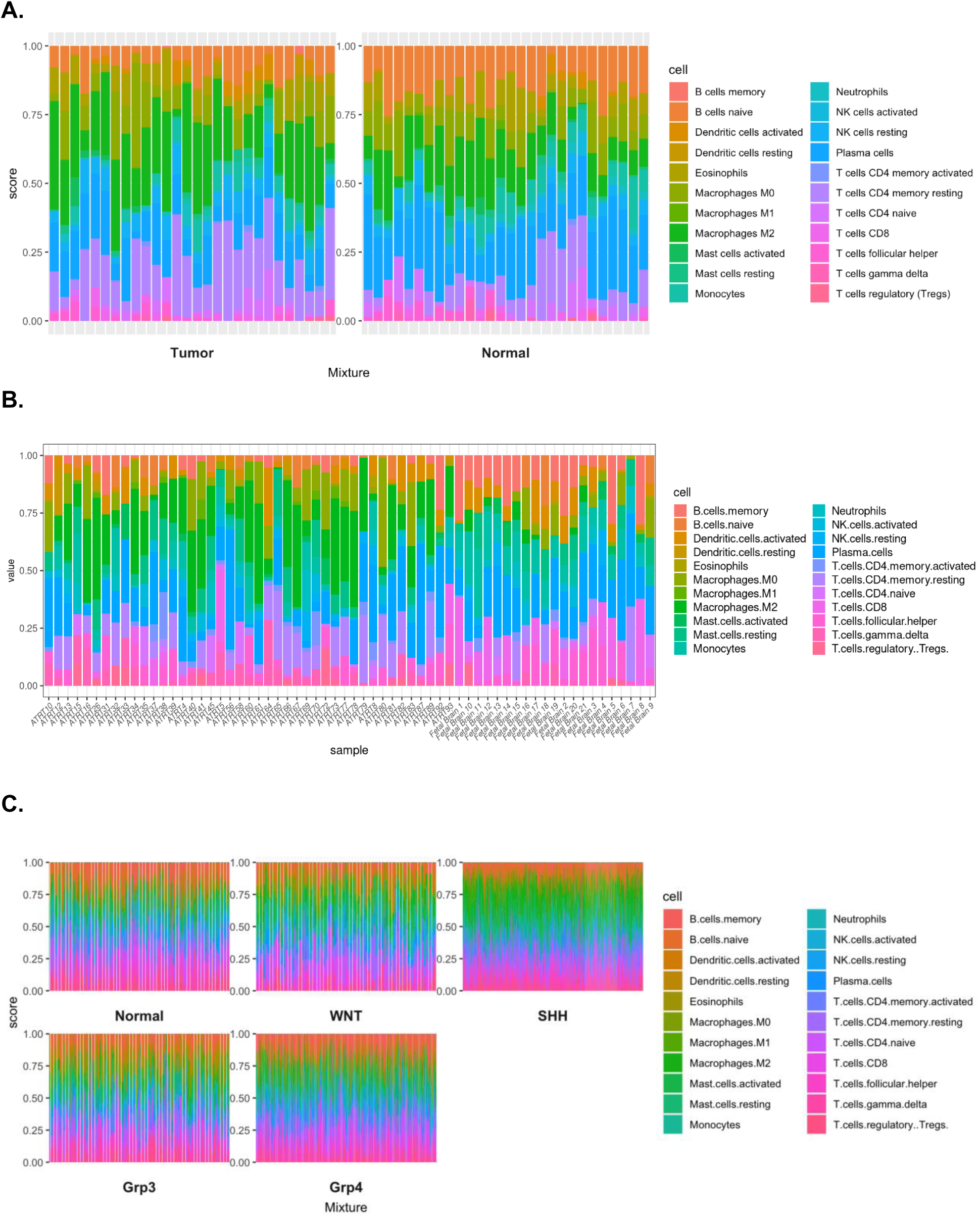
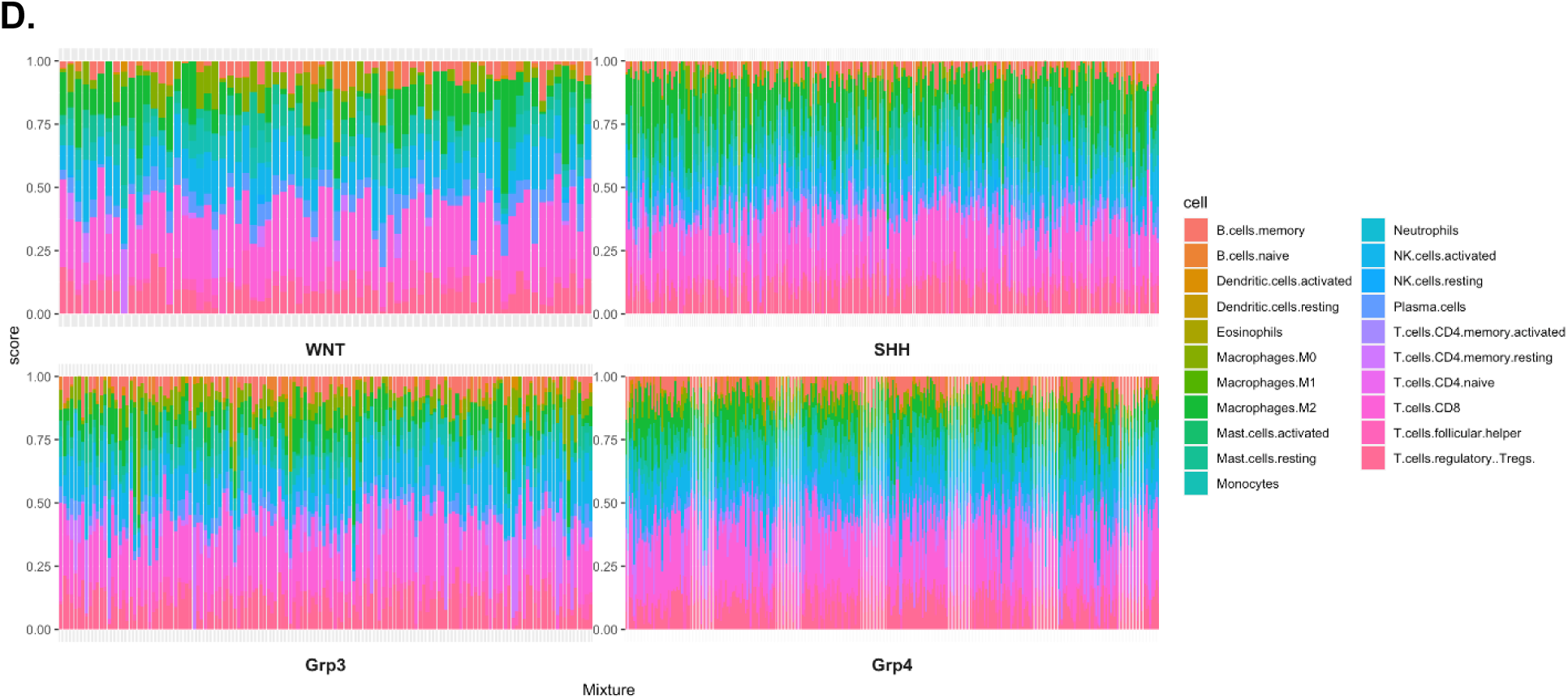
CIBERSORT for cell fractions in pediatric brain tumor samples. Gene expression data from various cohorts was input into CIBERSORTx (https://cibersortx.stanford.edu) where the abundance of various cell populations was estimated. (A) DIPG cohort, (B) ATRT cohort, (C) Medulloblastoma cohort, Swartling, and (D) Medulloblastoma cohort, Cavalli.

**S6.**
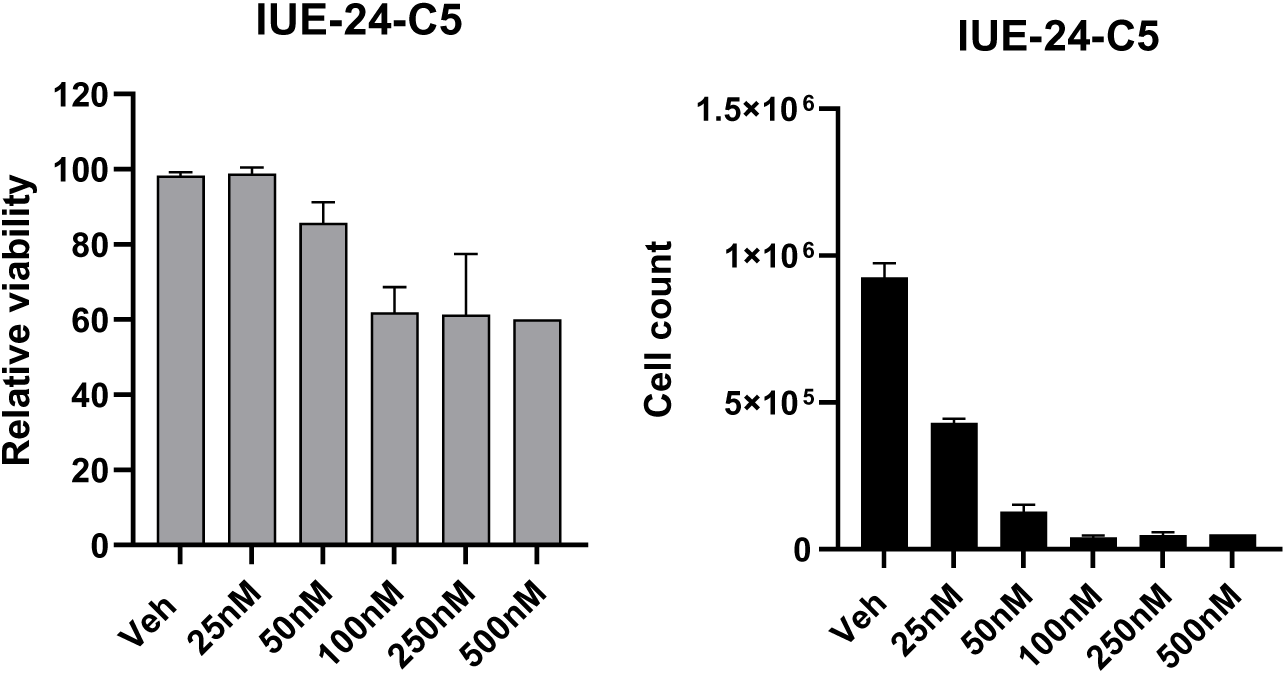
Treatment of IUE-24-C5 GEMM line with DAC for 7 Days. IUE-24-DIPG cells were treated with different concentrations (25nM, 50nM, 100nM, 250nM, 500nM) of decitabine for 7 days. Viability and total cell counts were determined (n=2).

**S7.**
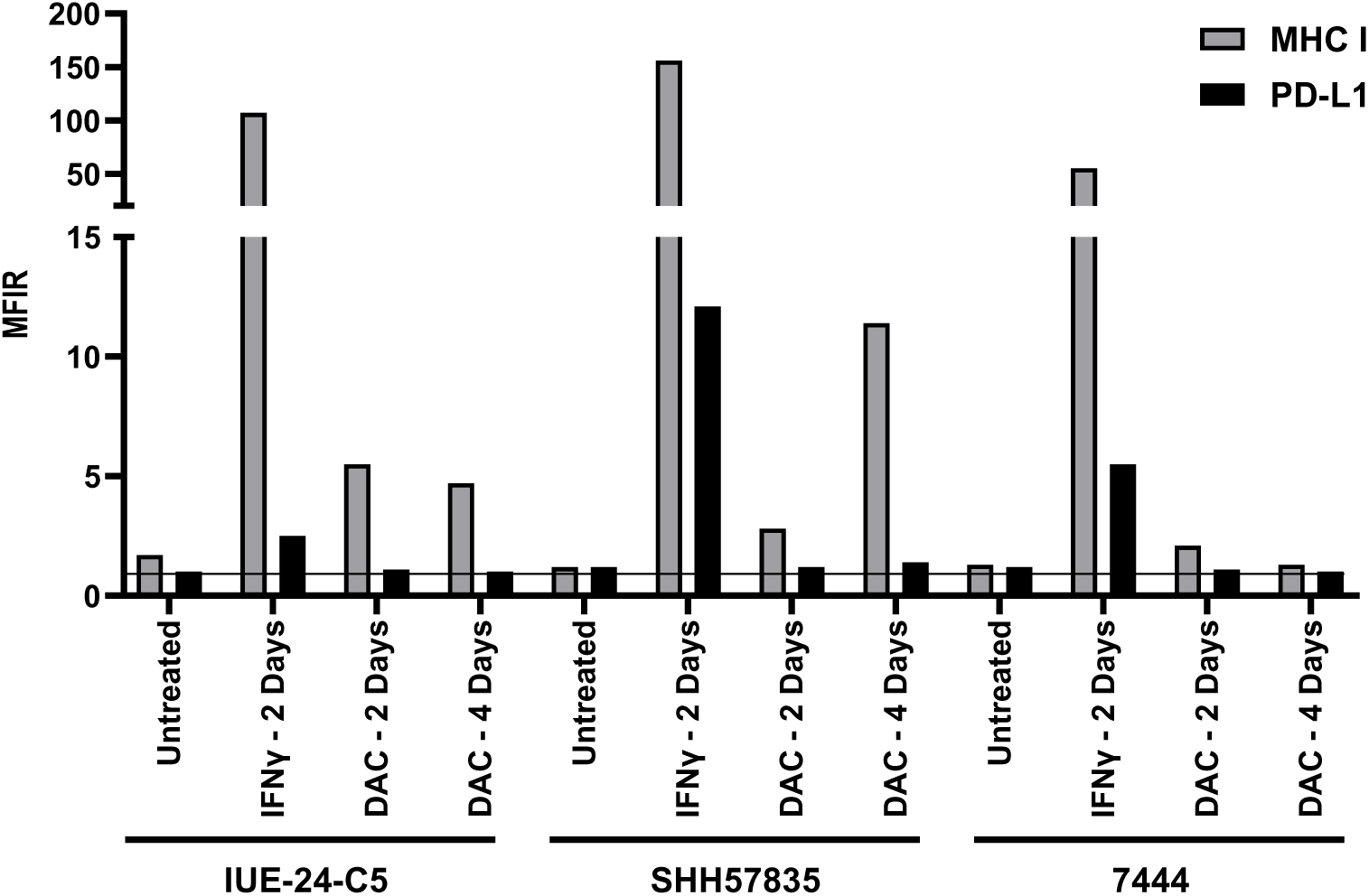
DNA methyltransferase inhibitor DAC induces MHC I expression in mouse syngeneic brain tumor cell lines. Baseline expression of MHC I and PD-L1 in IUE-24-C5, SHH57835, and 7444 mouse brain tumor cell lines (Untreated). Induction of MHC I expression following 2- or 4-day treatment of 0.5 μM DAC. IFNγ (50 ng/ml for 48 hours) was used as a positive control. MFIR equals the median fluorescence intensity of stained cells divided by the isotype control. Horizontal line indicates an MFIR of 1.

**S8.**
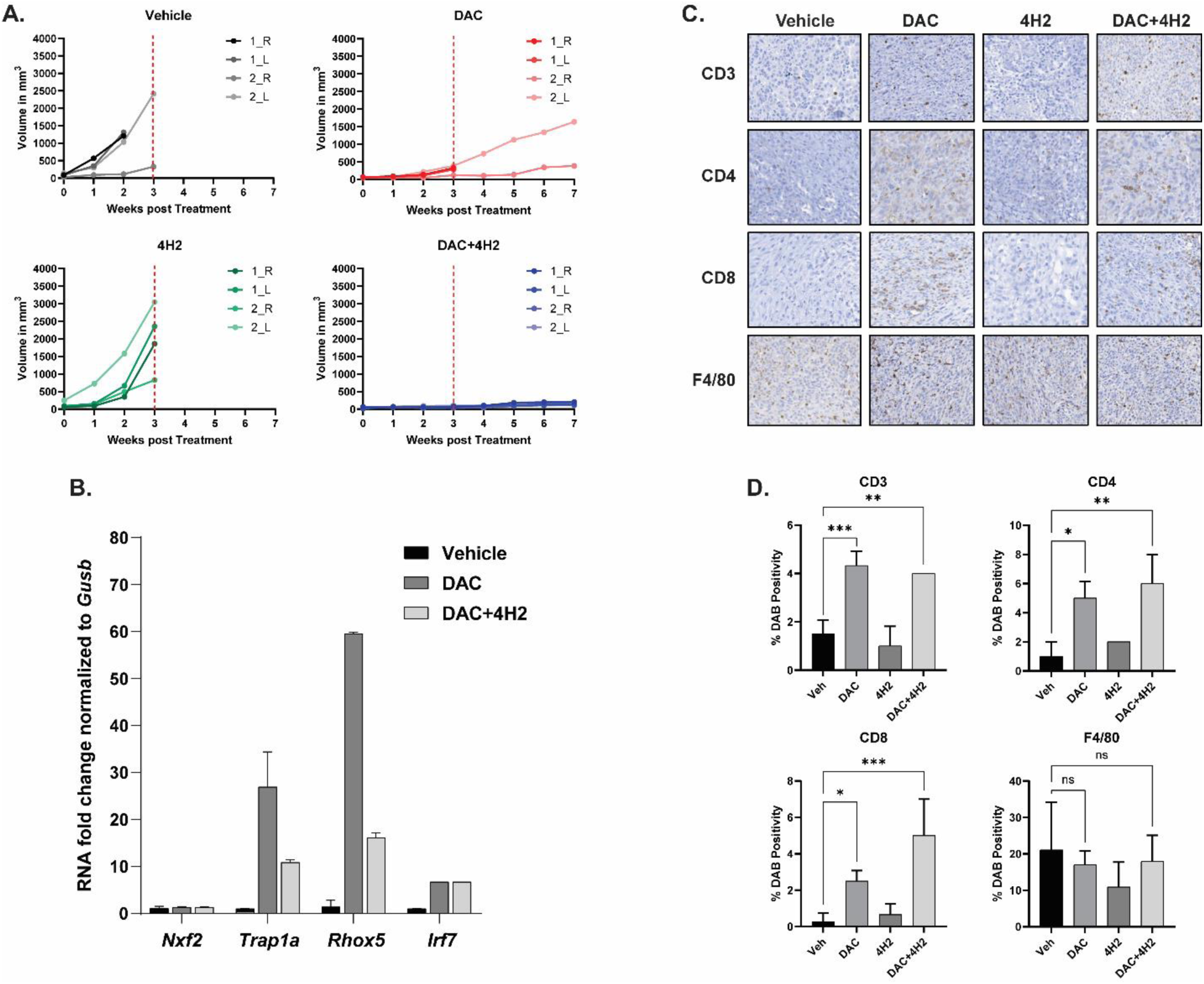
DAC in combination with 4H2 showed reduced tumor growth in the IUE-24-C5 syngeneic flank model. (A) Tumor cells were implanted in both right and left flanks of C57BL/6J mice and tumor volume was calculated by caliper measurements weekly. Changes in tumor growth were analyzed after 3 weeks of treatment (vertical red hatched line, when all vehicle treated mice succumbed to tumor burden). DAC and DAC+4H2 treatment showed significantly smaller tumors compared to vehicle and 4H2 treatment alone. (B) Gene expression analysis of flank tumors demonstrated that DAC treatment alone or in combination with 4H2 can induce *Trap1a* and *Rhox5* neoantigen expression along with *Irf7*, a transcription factor that regulates the expression of type I interferon genes. (C) IHC to identify the presence of various immune cell infiltrates in the flank tumors. (D) Quantitation of immune cell infiltrates as determined by percent DAB positivity.

**S9.**
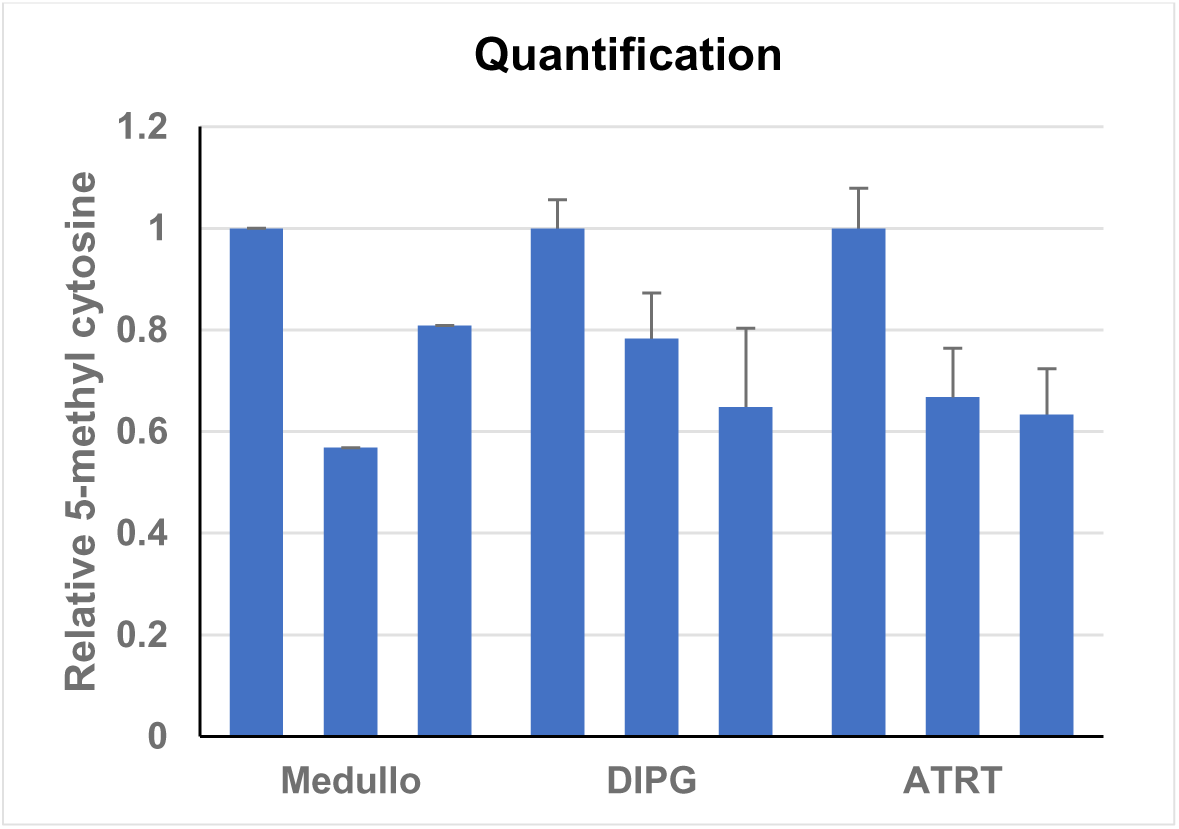
5-methylcytosine-specific dot blot assay was used to evaluate activity of DAC in mouse brain tumor tissue. Density of dot blots were measured and plotted as bar graphs.

**S10.**
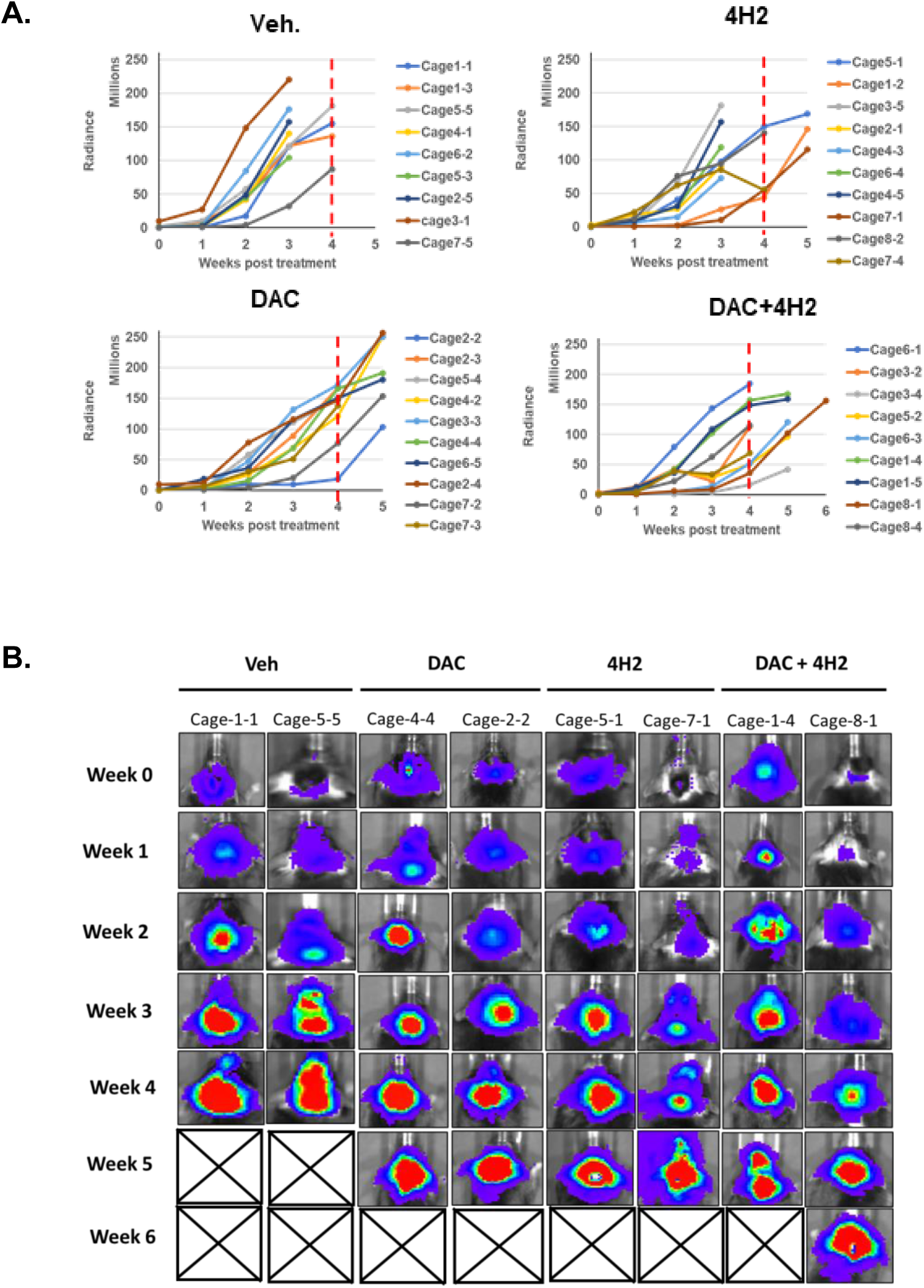
DAC and its combination with 4H2 showed inhibition of tumor growth in DIPG orthotopic model. (A) Luminescence of tumors over time as determined by IVIS imaging. DAC and DAC+4H2 treatment groups showed significantly improved survival (p<0.0001 and 0.0170, respectively compared to vehicle). Red vertical hatched line indicates 4-week post treatment commencement when all vehicle treated mice succumbed to tumor burden. (B) Representative IVIS images of mice from each treatment group.

**S11.**
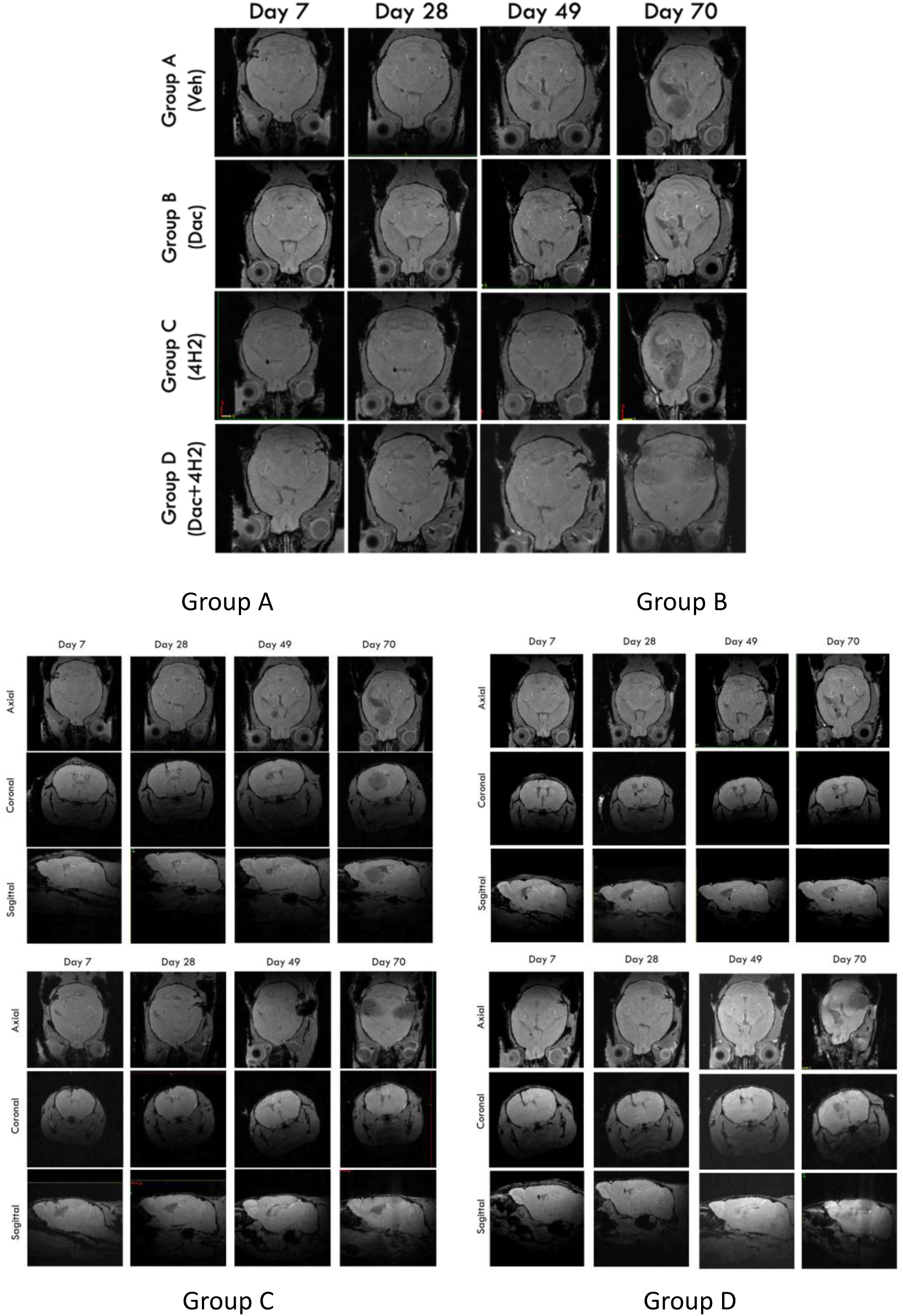
MRI Monitoring of ATRT tumors. Representative scans (axial, coronal, sagittal) taken at various time points during the study.

**S12.**
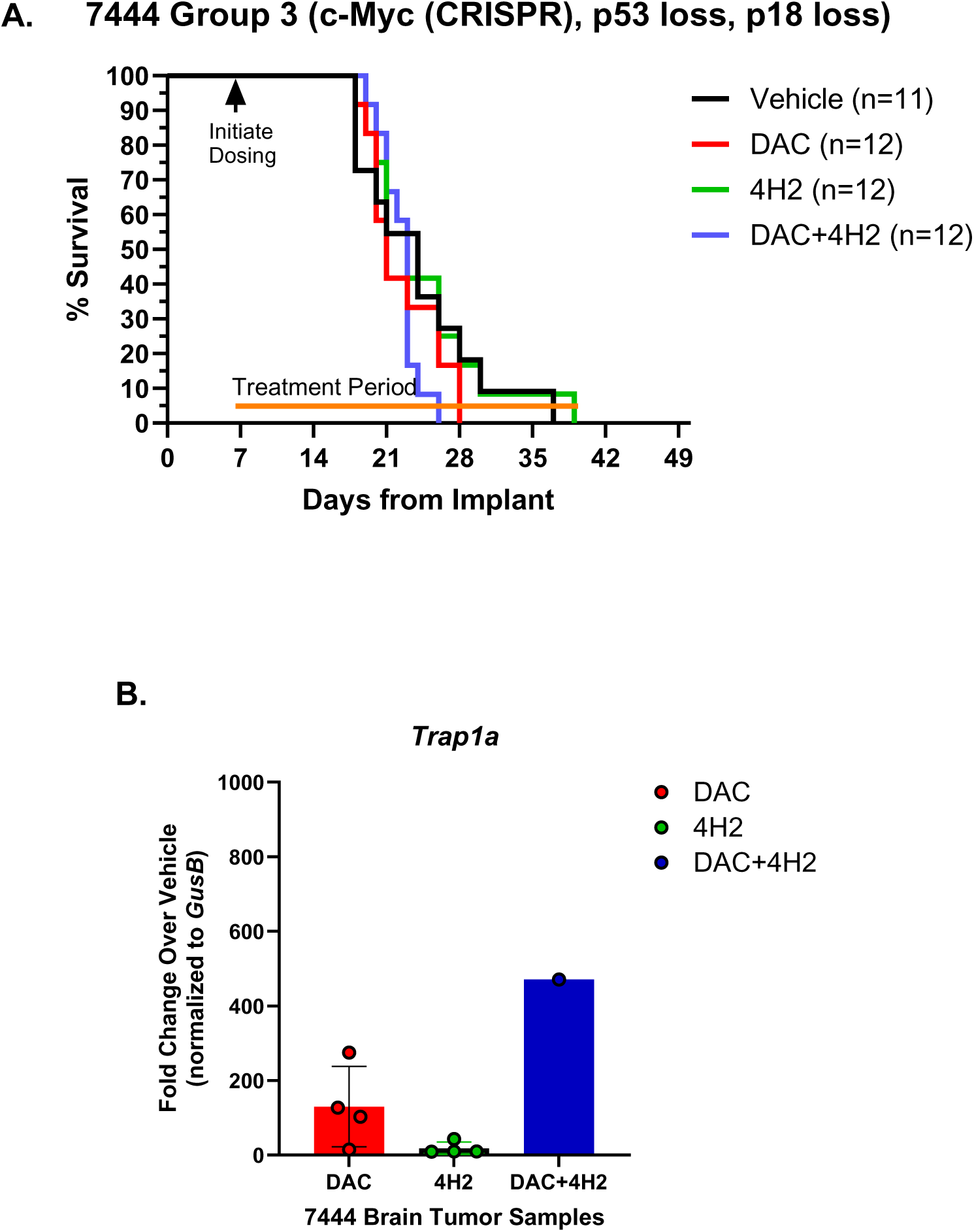
7444 Model Survival Curve and Neoantigen expression in tumor tissue. (A) Kaplan-Meier curves representing survival data for 7444 medulloblastoma model. (B) Evaluation of *Trap1a* neoantigen gene expression using quantitative PCR in tumor tissue harvested from 7444 syngeneic mice treated with either DAC, 4H2, or DAC+4H2.

**S13.**
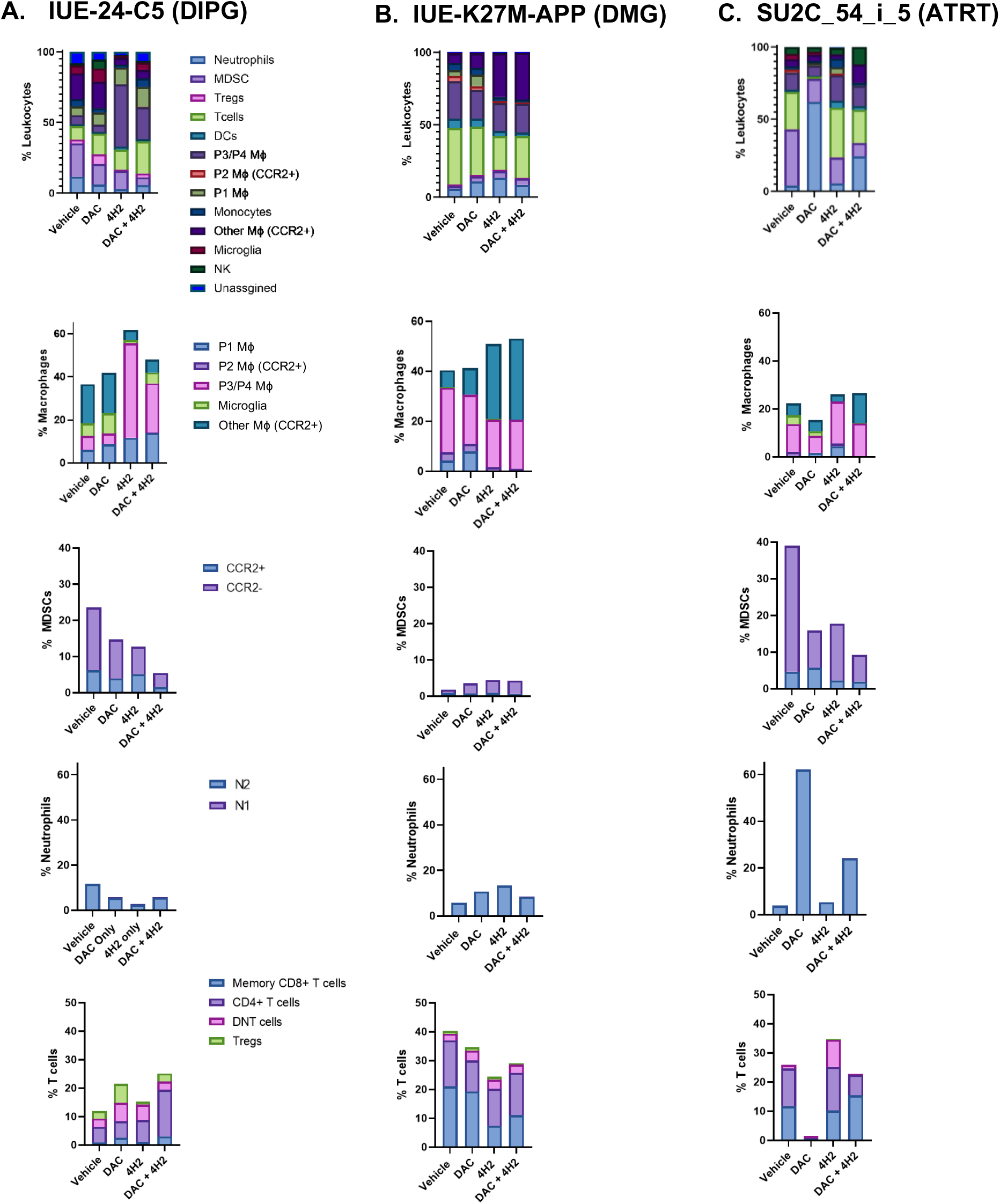
Additional CyTOF Data. Relative percentage of leukocytes, macrophages, MDSCs, neutrophils, and T cells in brain tumor tissue collected from vehicle, DAC, 4H2, and DAC+4H2 treated mice.

